# Genetic control of the error-prone repair of a chromosomal double-strand break with 5’ overhangs in yeast

**DOI:** 10.1101/2023.05.04.539391

**Authors:** Samantha Shaltz, Sue Jinks-Robertson

## Abstract

A targeted double-strand break introduced into the genome of *Saccharomyces cerevisiae* is repaired by the relatively error-prone nonhomologous-end joining (NHEJ) pathway when homologous recombination is not an option. A ZFN cleavage site was inserted out-of-frame into the *LYS2* locus of a haploid yeast strain to study the genetic control of NHEJ when the ends contain 5′ overhangs. Repair events that destroyed the cleavage site were identified either as Lys^+^ colonies on selective medium or as surviving colonies on rich medium. Junction sequences in Lys^+^ events solely reflected NHEJ and were influenced by the nuclease activity of Mre11 as well as by the presence/absence of the NHEJ-specific polymerase Pol4 and the translesion-synthesis DNA polymerases Pol ζ and Pol 11. Although most NHEJ events were dependent on Pol4, a 29-bp deletion with endpoints in 3-bp repeats was an exception. The Pol4-independent deletion required TLS polymerases as well as the exonuclease activity of the replicative Pol DNA polymerase. Survivors were equally split between NHEJ events and 1 kb or 11 kb deletions that reflected microhomology-mediated end joining (MMEJ). MMEJ events required the processive resection activity of Exo1/Sgs1, but there unexpectedly was no dependence on the Rad1-Rad10 endonuclease for the removal of presumptive 3′ tails. Finally, NHEJ was more efficient in non-growing than in growing cells and was most efficient in G0 cells. These studies provide novel insight into the flexibility and complexity of error-prone DSB repair in yeast.

## INTRODUCTION

DNA double-strand breaks (DSBs) are potentially toxic lesions that are repaired either by homologous recombination (HR), which uses an intact duplex as a template to restore the broken region, or by nonhomologous end joining (NHEJ), which directly ligates broken ends back together. In general, NHEJ is a more error-prone process, with out-of-register annealing between ends and/or end processing creating joints with small insertions or deletions. In addition, the rejoining of ends from different breaks can generate various types of genome rearrangements and has been implicated in recurrent oncogenic translocations. Although most DSBs are pathological and reflect either replication fork collapse, abortive topoisomerase reactions or DNA damage, programmed DSBs breaks are essential in some biological processes. During meiosis, for example, the Spo11 protein creates DSBs that initiate the HR necessary for creating genetic diversity and for ensuring proper chromosome segregation (KEENEY 2008). In mitosis, mating-type switching is initiated by the HO endonuclease in yeast (HABER 2012) and RAG proteins create DSBs that initiate the NHEJ-mediated joining of immunoglobulin gene segments in vertebrates (JUNG *et al*. 2006).

DSB repair pathways are highly conserved, and the yeast *Saccharomyces cerevisiae* has served as a model for defining relevant proteins and molecular mechanisms. Broken ends of all types are bound by the Ku (Ku70-Ku80) and MRX (Mre11-Rad50-Xrs2) complexes; both are absolutely required for NHEJ in yeast (DALEY *et al*. 2005b). Although not required for HR, MRX accelerates the initiation of 5′-end resection, which creates a 3′ tail that invades an homologous template and initiates HR (reviewed in SYMINGTON 2016; REGINATO AND CEJKA 2020). As part of the MRX complex, Mre11 nicks the 5′ terminated strand and subsequent 3′>5′ resection towards the break generates a free 3′ end. MRX activity is particularly important for eliminating end-attached proteins or terminal DNA damage and additionally removes Ku from ends to prevent NHEJ. In addition to its role in short-range resection, MRX facilitates the loading of long-range resection activities (Exo1 and Sgs1-Dna2) to promote efficient HR. HR requires a large suite of proteins to invade/copy the repair template as well as to resolve intermediates into final products (reviewed in SYMINGTON *et al*. 2014). In addition to Ku and MRX, NHEJ in yeast requires the dedicated Dnl4 DNA ligase (TEO AND JACKSON 1997) and the Pol4 DNA polymerase (WILSON AND LIEBER 1999).

Early NHEJ studies in yeast used transformation-based assays to assess the closure efficiency of linearized plasmids with defined end structures and to molecularly define the ligated products (reviewed in DALEY *et al*. 2005b). An advantage of this type of system is that end sequence can be manipulated *in vitro* to yield completely or partially complementary ends, or to generate completely incompatible ends (e.g., blunt ends or ends with different polarity).

With ends that have complementarity, the default is simple re-ligation. Following the annealing of ends with partial complementarity, gaps flanking the annealed region must be filled before ligation occurs. With 5′ overhangs, the filling of associated gaps is primed from a stably base-paired 3′ end. With 3′ overhangs, however, gap filling must occur from a 3′ end stabilized by at most a few base pairs and is more dependent on Pol4 (DALEY *et al*. 2005a). In contrast to 3′ overhangs, the recessed 3′ ends of 5′ overhangs can directly be extended in the absence of end annealing. With overhangs that lack complementarity or are incompatible, joining usually involves processing-uncovered microhomologies that flank the broken ends.

Mitotic studies of DSB repair in a chromosomal context have relied on endonucleases that create a single, targeted DSB and repair is monitored through selection of survivors or prototrophs. HO or I-*Sce*I, which generate breaks with 4-nt 3′ overhangs had been used extensively to study error-prone DSB repair (e.g. VILLARREAL *et al*. 2012; DENG *et al*. 2014). Zinc Finger Nucleases (ZFNs), which create 4-nt 5′ overhangs, and Cas9, which mostly creates blunt ends, have only rarely been used (LIANG *et al*. 2016; LEMOS *et al*. 2018; SHALTZ AND JINKS-ROBERTSON 2021). Because *S. cerevisiae* relies mainly on HR for the repair of genomic DSBs, NHEJ following endonuclease cleavage is studied either in the absence of a repair template or by disabling recombination. Precise rejoining of the ends by the NHEJ machinery regenerates the cleavage site, resulting in repetitive cycles of cleavage-ligation until a rare error-prone event renders the target sequence refractory to cleavage. Error-prone NHEJ typically involves minor addition or deletion of sequence from the broken ends and often reflects the annealing of small microhomologies within the overhangs, although microhomology is not a requirement. The overhang sequence dictates the spectrum of insertions/deletions and this sequence cannot be varied when DSBs are initiated with I-*Sce*I or HO. By contrast, use of a ZFN to create a DSB allows manipulation of the 5′ overhang sequence (LIANG *et al*. 2016; SHALTZ AND JINKS-ROBERTSON 2021). Reflecting the different reactions that can occur at 5′ versus 3′ overhangs, the kinetics and fidelity of repair also differ (LIANG *et al*. 2016).

In addition to the classical NHEJ pathway, yeast has an alternative end-joining pathway that is referred to as microhomology-mediated end joining (MMEJ). MMEJ is characterized by its Ku independence and a requirement for 6-14 bp of microhomology (reviewed in SFEIR AND SYMINGTON 2015). Finally, single-strand annealing (SSA) requires more extensive homology between direct repeats and is usually considered a variant of HR. In contrast to the canonical HR pathway, however, SSA is independent of the Rad51 strand-invasion protein (IVANOV *et al*. 1996), as is MMEJ (LEE AND LEE 2007). The transition from MMEJ to SSA occurs when the microhomology reaches 15-20 bp and SSA, but not MMEJ, has strong dependency on the Rad52 strand-annealing protein (VILLARREAL *et al*. 2012). It should be noted that higher eukaryotes lack a yeast-like MMEJ pathway and instead have an alternative end-joining pathway that is mediated by the Pol theta DNA polymerase (SFEIR AND SYMINGTON 2015).

We previously described a system that used a galactose-induced ZFN to create a site-specific DSB in the yeast *LYS2* gene (SHALTZ AND JINKS-ROBERTSON 2021). Insertion of an out-of-frame cleavage site allowed either the selection of NHEJ-mediated repair events that restored *LYS2* function or of repair events that simply allowed survival. Approximately half of the latter were Ku-independent MMEJ events (SHALTZ AND JINKS-ROBERTSON 2021). In the current study, this system was used to explore the genetic control of NHEJ- and MMEJ-mediated repair events.

## MATERIALS AND METHODS

### Media and Growth Conditions

All growth of yeast strains was at 30°C. For ZFN induction, cultures were grown non-selectively in YEP (1% yeast extract, 2% Bacto-peptone, 300 mg/liter adenine) supplemented with 2% raffinose (YEPR). Continuous ZFN expression was achieved by plating appropriate dilutions onto non-selective YEPGal, which contained 2% galactose, or onto selective SGal-lys synthetic medium (1.7 g/liter yeast nitrogen base, 0.5% ammonium sulfate, 2% agar, 2% galactose; all amino acids and bases except lysine). The total number of cells at the time of plating on galactose medium was determined by plating an appropriate dilution on YEPD medium (YEP plus 2% dextrose). Following transformation during strain constructions, selection was on YEPD containing the relevant drug or on synthetic medium missing the appropriate amino acid or base. Ura^-^ derivatives during two-step allele replacement were selected on synthetic medium supplemented with 5-fluoroorotic acid. Phenotypes following tetrad dissections were determined by replica-plating onto appropriate media. All mutant genotypes were confirmed by PCR.

### Strain constructions

All strains used were derived from the W303 background (*leu2-3,112 his3-11,15 trp1-1 ura3 ade2-1 CAN1 RAD5*) either by transformation or by mating and tetrad dissection. Each strain contained a galactose-inducible ZFN designed to cleave the Drosophila *rosy* locus (BEUMER *et al*. 2006) and a *lys2* frameshift allele containing a ZFN cleavage site. Genes encoding the ZFN constituent proteins RyA and RyB were integrated into the yeast as previously described (SHALTZ AND JINKS-ROBERTSON 2021) and ZFN cleavage sites were introduced into the *LYS2* locus using the *delitto perfetto* method (STORICI AND RESNICK 2006). Gene-deletion derivatives of SJR4848 were derived by one-step allele replacement using a plasmid-derived PCR cassette containing a selectable marker. The selectable cassettes used were *loxP-hphMX-loxP* from pSR955 (CHO AND JINKS-ROBERTSON 2019), *loxP-natMX-loxP* from pAG25 (GOLDSTEIN AND MCCUSKER 1999), *loxP-URA3Kl-loxP* from pUG72 and *loxP-TRP1-loxP* from pSR954 (CHO AND JINKS-ROBERTSON 2019). The *pol4-D367E* or *rev3-D975A* allele was introduced using *delitto perfetto*; the *mre11-D56N* or *pol3-DV* allele was introduced by two-step allele replacement using plasmid pSM444 (LLORENTE AND SYMINGTON 2004) or pY19 (JIN *et al*. 2001), respectively.

### Mutation Frequencies and Spectra

Independent cultures were grown in YEPR to an optical density (OD) reacofhed 0.3-0.6, at which point appropriate dilutions of “growing” cells were plated on YEPD to determine total cell number, on YEPGal to select survivors of continuous ZFN expression or on SGal-lys to select Lys^+^ revertants. To obtain a non-growing population of cells, incubation was continued for an addition day. G0 cells were isolated following the incubation of 25 ml YEPR cultures for seven days; small, unbudded G0 cells were obtained from the supernatant following low-speed centrifugation (KOZMIN AND JINKS-ROBERTSON 2013). The 95% confidence interval (CI) of the Lys^+^ frequency in each background was calculated using the standard error of the mean. Only “corrected” survival frequencies, which correspond to error-prone repair following continuous ZFN expression, are reported here and each was calculated by multiplying mean survival frequency on YEPGal by the fraction of sequenced colonies in the corresponding spectrum that had lost the ZFN cleavage site. The 95% CI for each corrected survival frequency was calculated by combining the 95% confidence interval for the measured frequency with the 95% CI for the proportion of sequenced colonies that had lost the ZFN cleavage site (MOORE *et al*. 2018). Frequencies of revertants and survivors in all backgrounds analyzed are in Table S1.

Mutation spectra in revertants and survivors were obtained by sequencing (Sanger method) a PCR-generated fragment spanning the ZFN recognition site. PCR failure was diagnostic of a large deletion that removed one or both of the primer-binding sites. The identity of the deletion was confirmed using a different set of primers, followed by sequencing the product (SHALTZ AND JINKS-ROBERTSON 2021). Complete spectra in revertants and survivors in all genetic backgrounds analyzed are in Table S2 and Table S3, respectively. The distributions of mutation types in different genetic backgrounds were compared using a global Chi-square test (Vassarstats.net). When a significant P value was obtained (p<0.05) comparisons of individual mutation types were then done and the P value for significance was adjusted by dividing 0.05 by the total number of comparisons performed (Bonferroni correction).

## RESULTS AND DISCUSSION

A galactose-regulated, heterodimeric ZFN designed to cleave the Drosophila *rosy* locus (BEUMER *et al*. 2006) was used to introduce a site-specific DSB in the *LYS2* gene (SHALTZ AND JINKS-ROBERTSON 2021). Each ZFN subunit contained three zinc fingers and recognized a 9-bp target sequence flanking a spacer of sequence 5′-ACGAAT (Figure 1A). Insertion of the 24-bp *rosy* target into *LYS2* created a -1 frameshift allele and net +1 mutations that restored the correct reading frame were selected by plating exponentially growing cultures on lysine-deficient medium containing galactose. Selection of surviving colonies on galactose-containing rich medium allowed a relatively unbiased assessment of events that eliminated the ZFN cleavage site. The genetic control of revertants, which reflected canonical NHEJ, and survivors, which reflected both NHEJ and MMEJ events, are described below.

**Figure 1.**
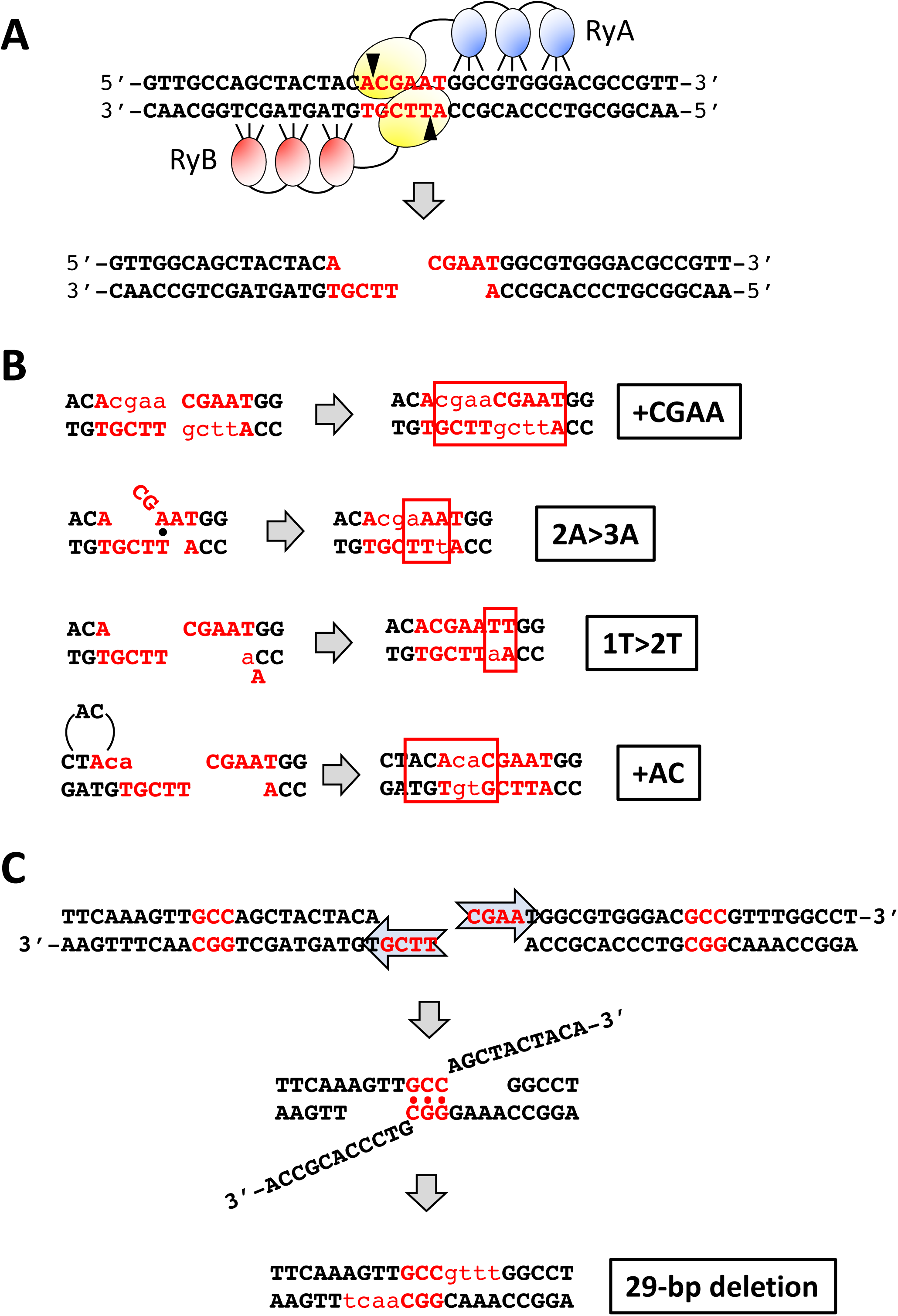
ZFN cleavage of spacer ACGAAT and error-prone repair events. (**A**) The sequence of the 24 bp *rosy* sequence inserted into *LYS2* is shown, with the three bases recognized by each zinc finger (blue or red ovals for RyA and RyB, respectively) indicated. The yellow ovals represent the dimerized *Fok*I domains; the 6-bp spacer is in red font and triangles indicate positions of enzyme-generated nicks that create 4-bp 5′ overhangs. (**B**) Sequences added during repair events are in lowercase red. Only the first three classes (+CGAA, 2A>3A and 1T>2T) generate Lys^+^ revertants; the fourth class (+AC) was only observed only among survivors. Complete filling of ends duplicates the region bounded by the ZFN nicks (+CGAA) while out-of-register pairing between an A and T in the overhangs generates the 2A>3A mutation. Duplication of the T (1T>2T) adjacent to the distal overhang can be generated by misincorporation-realignment, followed by ligation of the re-created 4-nt overhangs. The +AC event can be generated by a similar misincorporation-realignment mechanism during initial filling of the proximal overhang. (**C**) Mechanism for the NHEJ-dependent 29-bp deletion that is frequent among Lys^+^ revertants in the *pol4*1′ background. Resection of the 5′ ends allows pairing between GCC repeats (red) that flank the DSB. Subsequent removal of 3′ tails and filling of flanking gaps (red lowercase) completes repair.

### Effects of core NHEJ components on ZFN-induced Lys^+^ revertants

In a wild-type (WT) background, the frequency of Lys^+^ prototrophs was 2.03 x10^-4^ (Figure 2A) and there were three major classes of NHEJ events, each of which accounted for approximately 30% of revertants (Figure 2B; SHALTZ AND JINKS-ROBERTSON 2021). The first class (69/226) had a CGAA insertion (+CGAA) resulting from the complete fill in of each of the ZFN-generated 5′ overhangs (Figure 1B). The second major class (59/226) contained a 2A>3A expansion that is most easily explained by mis-annealing of the overhangs, followed by end trimming, gap filling and ligation. In the third class of Lys^+^ revertants (61/226) there was expansion to two thymines of a single thymine located immediately adjacent to the downstream ZFN-created overhang. We previously suggested that the 1T>2T event reflects misincorporation of an adenine, followed by realignment/slippage that regenerates the 4-nt overhang for direct ligation. Among the remaining events was a recurrent 29-bp deletion with endpoints in a GCC repeat (4/226; Figure 1C) as well as a variety of other minority events (33/226; see Table S2).

**Figure 2.**
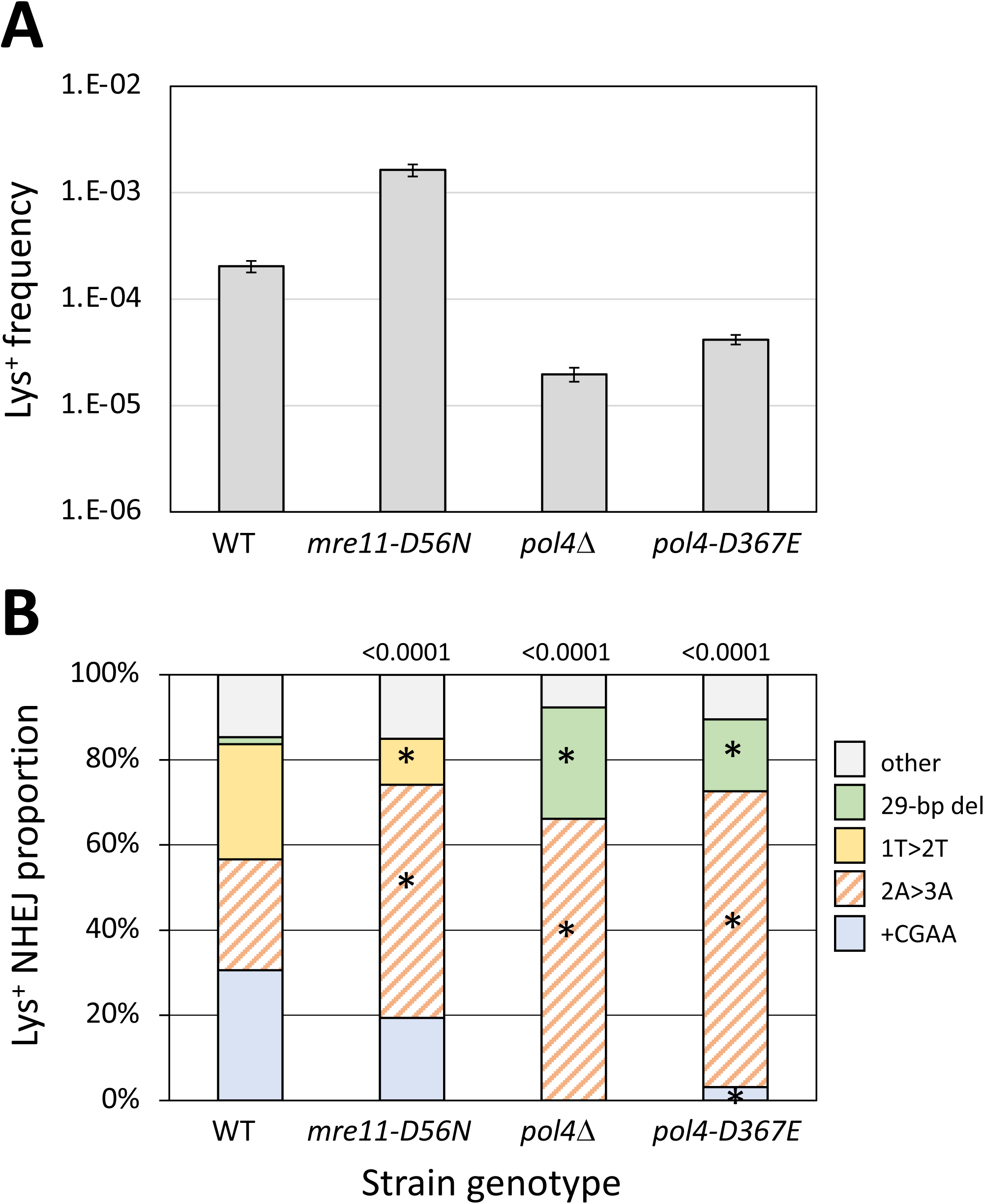
Frequencies and distributions of Lys^+^ revertants in strains with altered core NHEJ proteins. (**A**) Mean frequencies of Lys^+^ colonies; error bars are 95% confidence intervals for the mean. (**B**) Distributions of the five major NHEJ types among revertants. Overall distributions were compared to WT using a global 2 x 5 contingency chi square test (p values are above each spectrum) and if p<0.05, then individual mutation types were compared using the Bonferroni correction to determine significance (p<0.05/5); asterisks indicate a significant proportional class increase/decrease.

We previously reported that the frequency of Lys^+^ revertants decreased four orders of magnitude in a *yku70*1′ background (SHALTZ AND JINKS-ROBERTSON 2021) and we observed a similar reduction in *dnl4*1′ and *mre11*1′ strains (Table S1). Although Mre11 is required for NHEJ in yeast (MA *et al*. 2003), loss of only its nuclease activity stimulates NHEJ-mediated repair of 3′ overhangs, presumably by slowing initiation of the processive 5′ end resection required for HR (LEE AND LEE 2007; DENG *et al*. 2014). Introduction of the nuclease-dead *mre11-D56N* allele (MOREAU *et al*. 1999) into the ZFN system similarly stimulated Lys^+^ revertants 8-fold (Figure 2A) but additionally was associated with an unanticipated change in the spectrum of NHEJ events (Figure 2B; p<0.0001). Thus, in addition to a general repressive effect on NHEJ, Mre11 nuclease activity altered molecular outcomes.

In contrast to the complete absence of Lys^+^ prototrophs in *yku70*1′, *dnl4*1′ and *mre11*1′ backgrounds, there was only a 10-fold reduction in Lys^+^ frequency when *POL4* was deleted (Figure 2A), which is consistent with most, but not all, repair of HO-generated 3′ overhangs requiring Pol4 (LEE AND LEE 2007; TSENG *et al*. 2008). A similar Pol4 dependence of NHEJ was observed using plasmids with defined 3′ overhangs but, in contrast to the ZFN ends generated here in a chromosomal context, there was little or no Pol4 dependence with 5′ overhangs (DALEY *et al*. 2005a). Of the three major classes of Lys^+^ events observed following ZFN cleavage, only the 2A>3A event (121/183 revertants analyzed) persisted; there were no +CGAA or 1T>2T events detected (Figure 2B). A feature of the 2A>3A event that distinguishes it from the 1T>2T and +CGAA events is that it can occur through the annealing of 5′ tails, suggesting that Pol4 may be somewhat less important for gap-filling than for end-filling reactions. Of particular note, the 29-bp deletion event that was rare in WT (∼2%; 4/226) accounted for 26% (48/183) of ZFN-initiated events in the *pol4*1′ background. When converted to frequencies, Pol4 loss reduced the 2A>3A frequency 5-fold but had no effect on the 29-bp deletion. Finally, the relative importance of Pol4 presence versus its polymerase activity during NHEJ was examined using the *pol4-D367E* allele, which encodes a catalytically inactive protein (WILSON AND LIEBER 1999). While the Lys^+^ spectrum in the *pol4-D367E* background was identical to that in the *pol4*1′ mutant, the frequency of Lys^+^ revertants was 2-fold higher than in the *pol4*1′ strain. This suggests a minor structural role for Pol4 not previously detected (WILSON AND LIEBER 1999), which could reflect the stimulation of Dnl4-mediated ligation by Pol4 reported *in vitro* (TSENG AND TOMKINSON 2002).

### Effects of additional proteins on NHEJ-mediated repair of a ZFN-initiated break

The persistence of Lys^+^ colonies in the *pol4*1′ mutant indicates involvement of other DNA polymerases during repair of 5′ overhangs. The Pol ε replicative polymerase has been implicated, for example, in tail removal following the annealing of 3′ overhangs (TSENG *et al*. 2008). In the current study we focused on roles of the translesion synthesis (TLS) polymerases Pol ζ and Pol 11; *REV3* encodes the catalytic subunit of Pol ζ while the single subunit Pol 11 protein is encoded by *RAD30*. As observed following cleavage that generates 3′ overhangs (LEE AND LEE 2007), there was no significant change in the Lys^+^ frequency in the *rev3*1′ or *rad30*1′ single mutant. There was, however, a significant change in the proportional distribution of NHEJ events in each mutant (Figure 3). In the *rev3*1′ spectrum the proportion of +CGAA events increased 2-fold and accounted for 62% of events (58/94); the proportional decrease in 1T>2T events was significant (12/94) while that of 2A>3A events was not (14/94).

**Figure 3.**
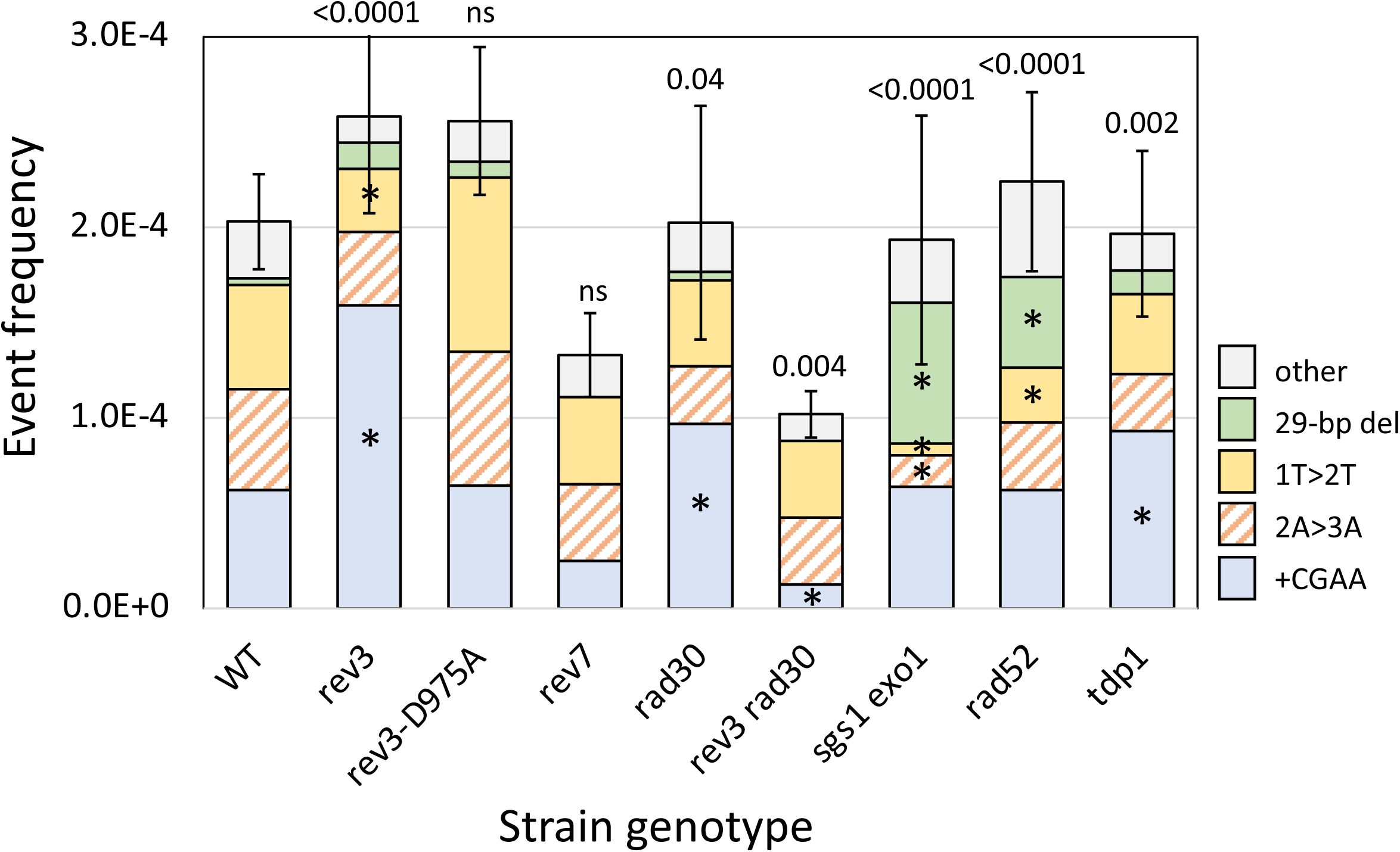
Altered frequencies and distributions of Lys^+^ revertants in mutant backgrounds. Mean frequencies of Lys^+^ colonies are shown; error bars are 95% confidence intervals for the means. The overall distribution of the five major NHEJ types in each mutant background was compared to WT using a 2 x 5 contingency chi square test (ns, not significant). If the global p value (above each spectrum) was less than 0.05, individual mutation classes were compared using the Bonferroni correction to determine significance (p<0.05/5); asterisks indicate a significant proportional increase/decrease.

Rev3 associates with chromatin near an HO-generated DSB (HIRANO AND SUGIMOTO 2006) and we used the *rev3-D975A* allele (JOHNSON *et al*. 2012) to determine whether the alteration in the NHEJ outcomes in the *rev3*1′ background reflects the presence of the protein or requires its catalytic activity. Both the frequency and spectrum of Lys^+^ colonies in the *rev3-D975A* catalytic mutant (JOHNSON *et al*. 2012) were indistinguishable from those of the WT parent, indicating that it is the presence of Rev3 that affects NHEJ outcomes. Rev7 interacts with Rev3 as part of the Pol ζ holoenzyme and we also examined its role in NHEJ. The effect of *REV7* deletion was distinct from that of *REV3* loss. Whereas the spectrum but not the frequency of Lys^+^ revertants was altered in the *rev3*1′ background, there was a slight reduction in frequency but the spectrum was unchanged in the *rev7*1′ mutant. The human equivalent of Rev7 (REV7, also known as MAD2L2) restrains end resection to limit HR and promote NHEJ (XU *et al*. 2015). Although a similar role for the yeast Rev7 has not been reported, the reduced NHEJ frequency in the *rev7*1′ background is suggestive of more robust resection in its absence.

In the *rad30*1′ background, the frequency of Lys^+^ revertants was not altered but there again was a significant change in the spectrum of NHEJ events (p=0.019). As in the *rev3*1′ background, the proportion of +CGAA events was elevated (45/94; p=0.0047); the proportion of the 2A>3A or 1T>2T events was not significantly altered (p>0.0125). Whereas individual deletion of *REV3* or *RAD30* affected the spectrum but not the frequency of Lys^+^ revertants, simultaneous deletion of both was associated with a 2-fold reduction in Lys^+^ frequency as well as a change in the NHEJ spectrum (p=0.0027). Instead of the proportional *increase* in +CGAA events observed in the single mutants, there was a specific *decrease* of this specific class in the *rev3*1′ *rad30*1′ double mutant. While the explanation for this is not obvious, especially given the *POL4* dependence of the +CGAA event, it underscores the complexity of interactions that take place during the error-prone repair of broken ends.

NHEJ-mediated repair in *rad52*1′ and *sgs1*1′ *exo1*1′ backgrounds was examined while measuring potential effects on MMEJ events in surviving colonies. The Rad52 protein is essential for homologous recombination and previous studies have reported either no (FRANK-VAILLANT AND MARCAND 2002; VILLARREAL *et al*. 2012) or very small changes (DENG *et al*. 2014) in NHEJ frequency in a *rad52*1′ background. Sgs1 and Exo1 are redundantly required for the processive resection of 5′ ends (MIMITOU AND SYMINGTON 2008) and their loss has no effect on NHEJ (VILLARREAL *et al*. 2012; DENG *et al*. 2014). While we similarly detected no significant change in the Lys^+^ frequency in either a *rad52*1′ or *sgs1*1′ *exo1*1′ background, the distribution of revertant types was altered in each (Figure 3). Particularly striking was the elevation in “other” NHEJ events: from 16% (37/226) in WT to 44% (41/94) in the *rad52*1′ and to 55% (52/94) in the *sgs1*1′ *exo1*1′ strain. In both backgrounds, this change reflected a significant increase in the Pol4-independent 29-bp deletion. A distinguishing feature of this specific NHEJ event is that it presumably requires at least some resection to expose complementary repeats, although it is limited by the processive resection of Sgs1/Exo1. If processive resection occurs normally, the data suggest that the 29-bp deletion may remain an option only if recombination cannot be initiated (*rad52*1′ background).

Yeast tyrosyl DNA phosphodiesterase (Tdp1) resolves the 5′- and 3′-phosphotyrosyl linkages associated with stabilized topoisomerases (POULIOT *et al*. 1999; NITISS *et al*. 2006) and has a 3′ nucleosidase activity that generates 3′-phosphate termini (INTERTHAL *et al*. 2005). Consistent with the latter activity, deletion of *TDP1* was associated with an increase in the complete fill-in of 5′ overhangs in a plasmid-based NHEJ assay (BAHMED *et al*. 2010). A similar effect was not observed, however, when repair of a ZFN-generated chromosomal break was examined by deep sequencing (LIANG *et al*. 2016). In our system *TDP1* deletion had no effect on the frequency of Lys^+^ revertants but did alter the spectrum of events (p=0.013). Among the three major classes of events there was a significant change only in the proportion of +CGAA events, (from 69/226 in WT to 53/112 in *tdp1*1′; p=0.0055). This is consistent with a subtle effect on Tdp1 on the ability to fill 5′ overhangs.

### Genetic control of the Pol4-independent 29-bp deletion

The model for the 29-bp deletion, which accounted for half of revertants in the *pol4*1′ background, requires 5′>3′ resection to expose GCC repeats that flank the ZFN cleavage site, annealing between complementary strands of GCC repeats, and cleavage of single-strand 3′ tails to allow filling of flanking gaps (Figure 1C). We first examined whether TLS polymerases are relevant to the Pol4-independent filling of the small gaps flanking the annealed segment. *REV3*, *RAD30* or both were deleted from the *pol4*1′ background and there were changes in the frequency and/or spectra of Lys^+^ revertants in each relative to the *pol4*1′ single mutant (Figure 4). The Lys^+^ frequency increased 2.0- and 3.7-fold, respectively, in the *pol4*1′ *rad30*1′ and *pol4*1′ *rev3*1′ backgrounds but was not significantly altered in the *pol4*1′ *rev3*1′ *rad30*1′ triple mutant.

**Figure 4.**
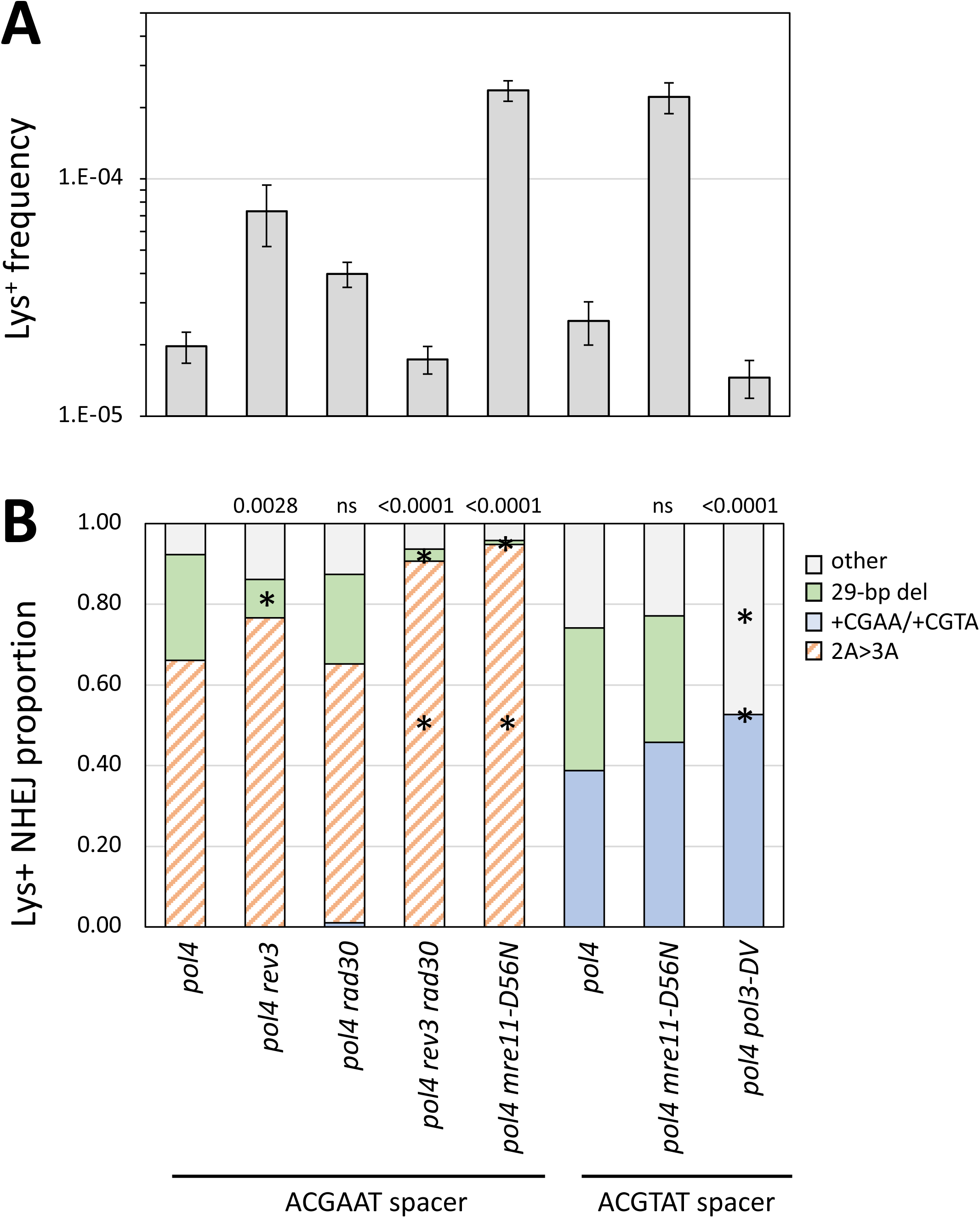
Genetic control of the Pol4-independent, 29-bp deletion. (**A**) Mean frequencies of Lys^+^ colonies; error bars are 95% confidence intervals for the means. (**B**) The overall distribution of the three major NHEJ types in each mutant background. With the ACGAAT spacer only the 2A>3A, the 29-bp deletion, and random “other” events were observed; there were no 1T>2T or +CGAA events. Changing the ACGAAT spacer to ACGTAT eliminated the 2A>3A event (orange cross-hatched), which was replaced by +CGTA (blue). Each distribution was compared to the corresponding WT using a 2 x 3 contingency chi square test (ns, not significant). If the global p value (above each spectrum) was less than 0.05, individual mutation classes were compared using the Bonferroni correction to determine significance (p < 0.05/3); asterisks indicate a significant proportional increase/decrease.

The proportion of the 29-bp deletion was reduced in the *pol4*1′ *rev3*1′ and *pol4*1′ *rev3*1′ *rad30*1′ backgrounds but was unaffected in the *pol4*1′ *rad30*1′ mutant. The frequencies of the 29-bp deletion were estimated by multiplying Lys^+^ frequencies and corresponding proportions of the 29-bp deletion. The 29-bp deletion frequency was unaffected by deletion of *REV3* or *RAD30* individually in the *pol4*1′ background but was reduced 10-fold in the *pol4*1′ *rev3*1′ *rad30*1′ triple mutant. These data demonstrate that Pol ζ and Pol 11 have redundant roles during the Pol4-independent gap filling required to generate the 29-bp deletion.

Although sporadic deletions involving other short repeats were observed among revertants in all genetic backgrounds (Table S2), the GCC repeats at the endpoint of the 29-bp deletion were notable because they are close to and symmetrically flank the ZFN cleavage site. Furthermore, the frequency of the 29-bp deletion was elevated when the long-range resection pathways were eliminated (*exo1*1′ *sgs1*1′ double mutant; Figure 3). Given the repressive effect of resection on this event and the symmetry/proximity of the GCC repeats to the ZFN cleavage site, we considered the possibility that Mre11 nuclease activity might be required to expose the repeats within short 3′ overhangs. Mre11 nicks the 5′ strand 30-35 nt from a Ku-bound end and expansion of the nick into a gap by its 3′>5′ exonuclease activity degrades the 5′ end to displace Ku and create a short 3′ tail (reviewed in REGINATO AND CEJKA 2020). To examine the relevance Mre11 nuclease activity to the NHEJ-dependent 29-bp deletion, the nuclease-dead *mre11-D56N* allele (LLORENTE AND SYMINGTON 2004) was introduced into the *pol4*1′ background. The Lys^+^ frequency increased 12-fold in the double mutant relative to the *pol4*1′ single mutant and this was accompanied by a change in the distribution of revertant types (Figure 4). Almost all revertants had the 2A>3A event (91/96; p<0.0001) and only a single 29-bp deletion was observed. The *mre11-D56N* allele thus had a strong stimulatory effect on the frequency 2A>3A event, but its effect on the frequency of the 29-bp deletion was unclear.

To preclude occurrence of the 2A>3A event and allow more specific focus on the 29-bp deletion in the *pol4*1′ background, we changed the spacer sequence of the ZFN cleavage site from ACGAAT to ACGTAT. With the new ACGTAT spacer in the *POL4* background, there were similar numbers of +CGTA and 1T>2T events (43/116 and 63/116, respectively), as there were with the original spacer, and the 29-bp deletion remained rare (2/116). Given the absence of +4 and 1T>2Tevents with the original ACGAAT spacer in the *pol4*1′ background, we assumed that the 29-bp deletion would be present in almost all Lys^+^ colonies derived using the new ACGTAT spacer. In the *pol4*1′ background with the new spacer, however, half of revertants contained the +CGTA event (54/37) and 30% had the 29-bp deletion (37/118); there still were no 1T>2T events. In the absence of Pol4, the data suggest that the end-annealing that generates the 2A>3A event may precede complete filling and obscure it occurrence.

Elimination of Mre11 nuclease activity in a *POL4* background with the new spacer increased the frequency of Lys^+^ revertants 14-fold and altered the proportions of +CGTA and 1T>2T events (Tables S1 and S2). This mirrors the differential effects of the *mre11-D56N* allele on the NHEJ spectrum observed with the original ACGAAT spacer. Introduction of the *mre11-D56N* allele into the *pol4*1′ background resulted in a 9-fold increase in Lys^+^ frequency relative to the *pol4*1′ single mutant, which was similar to the increase observed with the original spacer. In contrast to the large proportional decrease in the 29-bp deletion in the *pol4*1′ *mre11-D56N* background with original spacer, however, there was no reduction with the new spacer. This demonstrates that Mre11 nuclease activity is not required to expose the GCC repeats at the endpoints of the 29-bp deletion.

In the model depicted in Figure 1C end resection allows annealing between complementary strands of the GCC repeat, thereby creating 3′ tails that must be removed prior to gap filling and ligation. The Rad1-Rad10 nuclease is required for the removal of long 3′ tails during SSA, but tails <30 nt are efficiently removed by the exonuclease activity of Pol δ (PAQUES AND HABER 1997). In the *pol4*1′ strain containing the new spacer sequence, elimination of the exonuclease activity of Pol δ (*pol3-DV* allele; (JIN *et al*. 2001) reduced the Lys^+^ frequency 2.5-fold and this reflected the complete absence of the 29-bp deletion among revertants (0/95 revertants; p<0.0001; Figure 4). These data demonstrate that the exonuclease activity of Pol 8 is required during creation of the 29-bp deletion.

The data presented above present a conundrum with respect to the genetic control of the NHEJ-dependent 29-bp deletion. It was antagonized by processive nuclease activity (*sgs1*1′ *exo1*1′ background) and yet did not appear to depend on the nuclease activity of Mre11 (*pol4*1′ *mre11-D56N* background). How then are the complementary strands of the GCC direct repeats exposed? One possibility is that the DNA melting activity of the MRX complex is responsible.

This requires the ATPase activity of Rad50 (CANNON *et al*. 2013), which also is required for NHEJ in yeast (ZHANG AND PAULL 2005). The strong dependence of the 29-bp deletion on the exonuclease activity of Pol δ (*pol4*1′ *pol3-DV* mutant) suggests an alternative possibility in which it is the degradation of the recessed 3′ ends by Pol δ that exposes the GCC repeats. Regardless of how the complementary strands of the GCC repeats are exposed, the Ku dependence of the 29-bp deletion suggests either that Ku remains associated with the ends or that Ku re-engages the ends after displacement. Ku interacts with duplexes with 30-nt tails almost as well as with fully duplex DNA *in vitro* (FALZON *et al*. 1993); longer, 60-nt tails are not efficiently bound by Ku (FOSTER *et al*. 2011).

### Survivors of continuous ZFN expression

Selection for colonies on rich medium containing galactose provides an unbiased analysis of end-joining events that confer resistance to continuous ZFN expression. As reported previously, only half of survivors (83/155) in the WT background lost the enzyme cleavage site (SHALTZ AND JINKS-ROBERTSON 2021) and these were of two major types: a Ku-dependent +AC event that expanded two copies an AC dinucleotide spanning the 6-bp ZFN spacer (36/83) and Ku-independent large deletions that removed the cleavage site (47/83). The +AC event was not detectable in the reversion assay, which selects net +1 events, and was hypothesized to arise by misincorporation-slippage during proximal end filling (Figure 1B). The rarity of net +1 events in the survivor assay indicates that they are minor events relative to +AC. The large deletions were either 1.2 kb (36/47) or 11.7 kb (11/47) in size (Figure 5A) and because the deletion junctions were in 13- or 14-bp direct repeats, respectively, we concluded that these were MMEJ events. In the current analyses, the frequencies and profiles of survivors were examined in the genetic backgrounds described above for Lys^+^ revertants. Only those mutants that were different from the WT strain in terms of the frequency and/or spectrum of site-loss survivors are discussed (see Tables S1 and S3 for all data).

**Figure 5.**
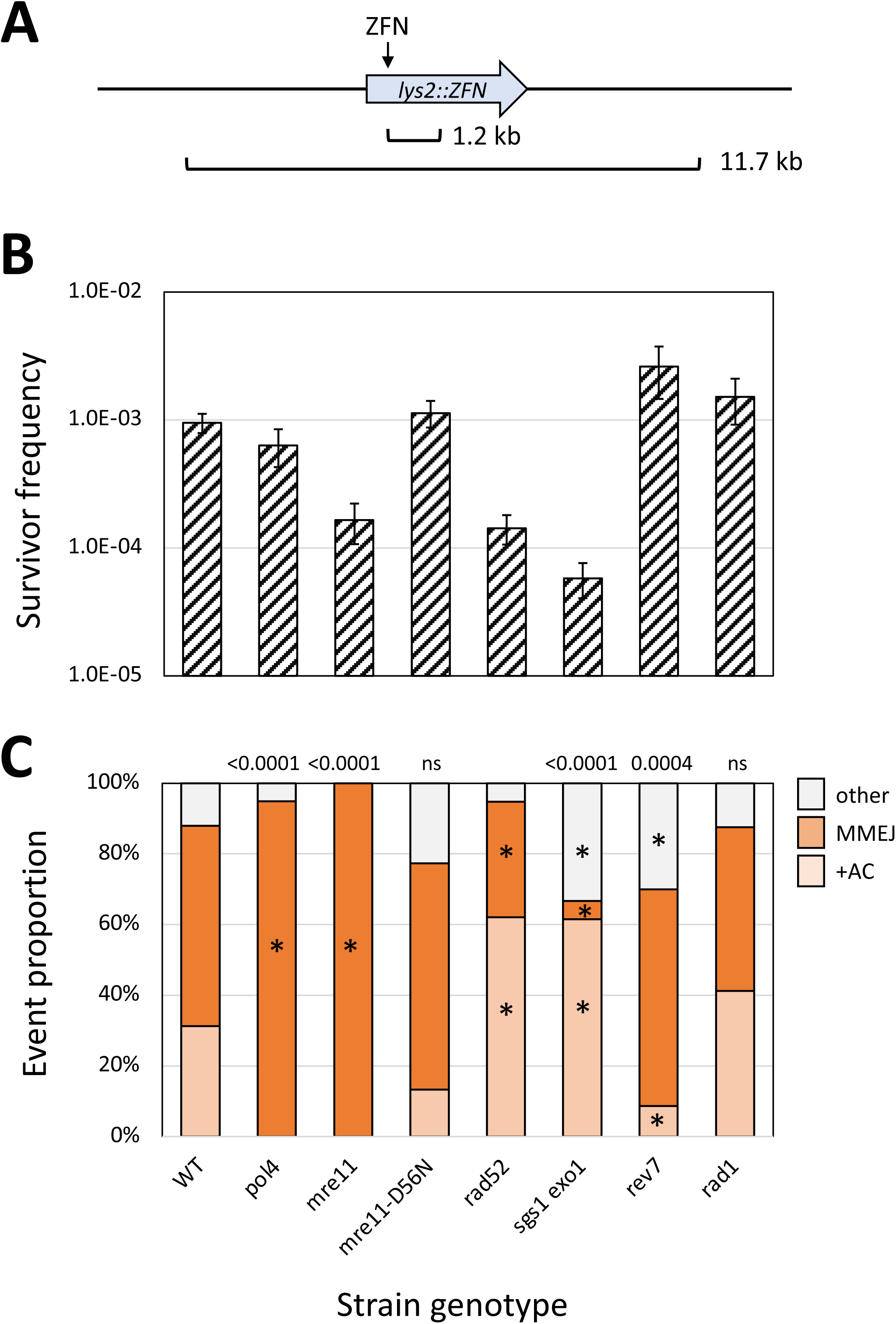
Genetic control of the survivor frequency and spectrum. (**A**) The two types of recurrent large deletions detected among surviving colonies. The 1.2 kb deletion as endpoints in 13-bp direct repeats (CCAAGCTACTACA), one of which overlaps the DNA binding site of the RyB subunit of the ZFN. The 11.7 kb deletion has endpoints in 14-bp direct repeats (TGGAAAAAAAAAAA) and removes the *LYS2*-flanking genes *TKL2* and *RAD16* (not shown). (**B**) Mean frequencies of surviving colonies that lost the ZFN cleavage site; error bars are 95% confidence intervals for the means. (**C**) The distributions of the three major survivor types are shown. Each mutant distribution was compared to the WT using a 2 x 3 contingency chi square test (ns, not significant). If the global p value (above each spectrum) was less than 0.05, individual mutation classes were compared using the Bonferroni correction to determine significance (p<0.05/3); asterisks indicate a significant proportional increase/decrease.

Because NHEJ and MMEJ proportions are similar among survivors, the complete loss of either pathway would be expected to reduce the survival frequency only 2-fold. Indeed, as reported previously in the *yku7*01′ background (SHALTZ AND JINKS-ROBERTSON 2021), the survival frequency was reduced 2-fold in the NHEJ-deficient *pol4*1′ (Figure 5B-C) and *dnl4*1′ (Tables S1 and S3) backgrounds and only large deletions were detected. Although NHEJ events were also absent in an *mre11*1′ strain, there was 5.9-fold reduction in overall survival frequency rather than the expected 2-fold (Figure 5B). Survivor frequencies were also low in *rad52*1′ single and *sgs1*1′ *exo1*1′ double-mutant backgrounds, with reductions of 6.7- and 16.7-fold, respectively, relative to WT. Impaired growth was a common feature of these three mutant backgrounds, and we suggest that this may be responsible for the large decrease in survivor frequency. We favor this interpretation because, in contrast to the survivor assay, the NHEJ frequency as measured by Lys^+^ reversion was not affected in either the *rad52*1′ or *sgs1*1′ *exo1*1′ background (Figure 3). It is possible that repetitive cleavage by a ZFN impairs viability more when cells can continue to divide (nonselective rich medium) than when cells are plated under conditions where error-prone repair must restore prototrophy before cells can begin dividing.

In addition to reduced survival, the spectrum of events among survivors was altered in the *rad52*1′ and *sgs1*1′ *exo1*1′ backgrounds relative to WT (p=0.0013 and p<0.0001, respectively). There was a reduction in the proportion of large deletions in the *rad52*1′ mutant, (from 47/83 to 19/58; p=0.009), suggestive of a partial requirement of Rad52 for the MMEJ-mediated large deletions. In an earlier study that systematically examined the effect of repeat size on microhomology-mediated end joining (VILLARREAL *et al*. 2012), Rad52 promoted end joining for repeats 15 bp or larger, suggesting a variation of SSA as the mechanism, and strongly inhibited end-joining between repeats 12 bp or smaller. The sizes of the repeats at the MMEJ endpoints in the current study (13-14 bp) are in the transition zone for Rad52 dependence. In the *sgs1*1′ *exo1*1′ double mutant the proportion of large deletions was reduced 10-fold: from 47/83 in WT to only 2/39. This reduction likely reflects the extensive resection required to expose the junction repeats of large deletions and is consistent with results obtained with similar large deletions (VILLARREAL *et al*. 2012). This is in contrast to the suppressive effect of Sgs1/Exo1 resection (as well as Mre11 presence) with respect to Ku-independent deletions between 12-bp repeats very close to I-*Sce*I-generated ends (DENG *et al*. 2014). The resection-related suppression is similar to that described above for the Pol4-independent 29-bp deletion, although the 29-bp deletion is Ku- and Mre11-dependent (see above).

Because the nuclease activity of Mre11 inhibits NHEJ, an elevation in the NHEJ-dependent +AC event was expected in an *mre11-D56N* background. There, however, was neither a change in survivor frequency nor spectrum relative to WT. This result provides further support for an influence of Mre11 nuclease activity on specific NHEJ outcomes, as inferred from the variable effect of the *mre11-D56N* allele on Lys^+^ revertant types in the WT and *pol4*1′ backgrounds. Finally, there was 2.7-fold increase in survivor frequency in a *rev7*1′ background that specifically reflected an increase in MMEJ; no effect on frequency or spectra was observed upon loss of *REV3* (Table S1). These data more strongly support a potentially suppressive role of Rev7 on processive resection in yeast, as inferred previously from the slightly increased frequency of Lys^+^ revertants.

The Rad1-Rad10 endonuclease is required to remove 3′ tails during SSA (IVANOV AND HABER 1995) and a similar importance for Rad1 in MMEJ was inferred through analysis of HO-initiated 2-kb deletions between 18-bp direct repeats (VILLARREAL *et al*. 2012). The strong Rad52 dependence for deletions between 18-bp, however, suggests that the events were a variation of SSA. We thus re-examined the requirement for Rad1 for the large, NHEJ-independent deletions in our system. With the smaller (13- and 14-bp) repeats, there was no significant change in either the frequency or the distribution of survivor types in a *rad1*1′ background (Figure 5; see Tables S1-S3 for similar *rad10*1′ data). To potentially identify the relevant structure-specific nuclease that removes 3′ tails during MMEJ, we examined survival in *mus81*1′ single and *rad1*1′ *mus81*1′ double mutants, as well as in an *slx4*1′ background. Mus81 can process 3′ flaps (SCHWARTZ AND HEYER 2011) while Slx4 is a scaffold for multiple structure-specific endonucleases (CUSSIOL *et al*. 2017). There was no proportional decrease in large deletions in any of these additional mutant backgrounds (Table S3). Either a different nuclease is relevant, or there is functional redundancy between the nucleases examined. Just as there is a size-related transition in Rad52 dependence for microhomology-mediated deletions (VILLARREAL *et al*. 2012), there may be a similar transition in terms of a requirement for Rad1-Rad10 in 3′-tail removal. One interesting possibility is that Rad52-driven annealing dictates whether subsequent tail removal is dependent on Rad1-Rad10.

### The physiological state of cells affects repair of ZFN-induced DSBs

NHEJ-mediated ligation of a linearized plasmid is more efficient when non-growing (NG) cells are transformed than when growing cells are transformed (KARATHANASIS AND WILSON 2002). To examine the effects of growth state on error-prone repair of a chromosomal DSB, either non-growing (NG) or isolated G0 cells were plated in the presence of galactose. The Lys^+^ frequency in NG cells was 2.4-fold higher than in growing cells and there were proportionally fewer +CGAA events among revertants (Figure 6A). When the event types were converted to frequencies, however, the +CGAA frequency was unaltered while the frequencies of 2A>3A, 1T>2T and other NHEJ events were each elevated about 3-fold. There was a further stimulation of the Lys^+^ frequency when G0 cells were plated: 6.8- and 2.8-fold relative to growing and NG cells, respectively. Interestingly, there was a very large proportional increase in the +CGAA class of events in G0 cells that corresponded to a 10-fold increase in its frequency relative to growing/NG cells; the frequencies of the other classes changed little relative to NG cells.

**Figure 6.**
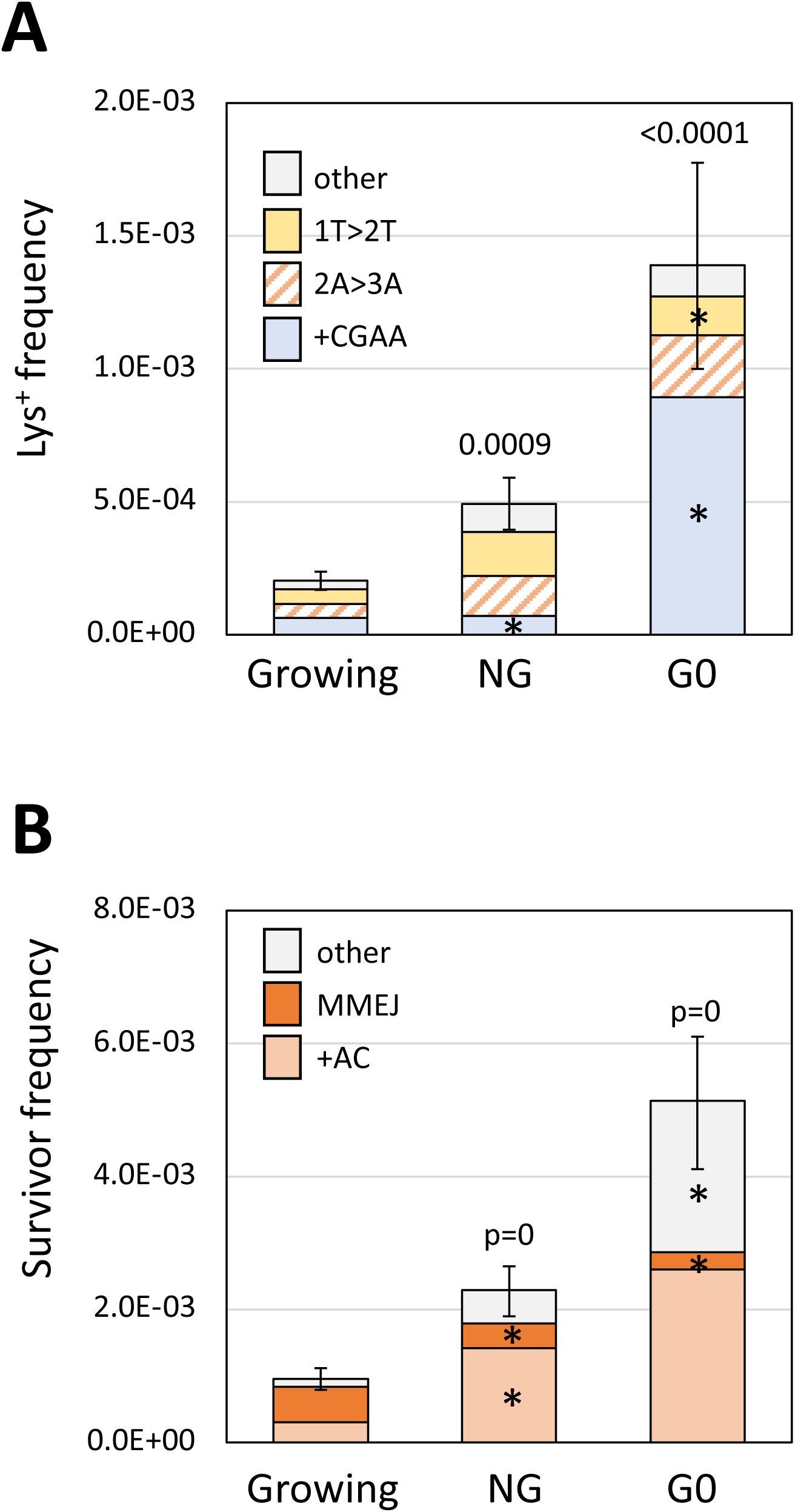
Alterations in error-prone DSB repair in NG and G0 cells. (**A**) Mean Lys^+^ frequencies and revertant-type distributions in growing, NG and G0 cells. (**B**) Mean survivor frequencies and event-type distributions in growing, NG and G0 cells. Error bars are 95% CIs for mean frequencies. Event-type distributions were compared to growing cells using a 2×4 or 2×3 contingency chi square test for revertants or survivors, respectively; ns, not significant. I the global p value was <0.05 (above each spectrum), then ndividual mutation types were compared using the Bonferroni correction to determine significance; asterisks indicate a significant proportional increase/decrease.

In terms of surviving colonies that had lost the ZFN cleavage site, the overall frequency also was elevated relative to growing cells: 2.4-fold in NG cells and 5.4-fold in G0 cells (Figure 6B). The spectrum of survivors was also altered. As expected of events that require extensive resection, the proportion of large deletions was greatly reduced: from 0.56 (47/83) in growing cells to 0.16 (15/92) and 0.05 (4/79) when NG and G0 cells, respectively, were plated. Given the large increase in survival, however, the large-deletion frequency was reduced only about 2-fold in NG and G0 cells. In terms of frequency, the +AC event was elevated 4.7-fold in NG cells and 8.7-fold in G0 cells; the frequency of “other” events was similarly elevated. The data demonstrate that NHEJ-mediated repair of a ZFN-generated DSB is more efficient in NG/G0 cells than in growing cells, and that physiological state additionally affects how the resulting ends are modified during error-prone repair.

## CONCLUSIONS

- Following ZFN cleavage, half of error-prone repair events reflected small insertions/deletions at the cleavage site (NHEJ) and half were large deletions (MMEJ).
- In contrast to the complete dependence of NHEJ on Ku, Mre11 and Dnl4, loss of Pol4 reduced NHEJ only 10-fold. Although most NHEJ events were dependent on Pol4, a recurrent 29-bp deletion was Pol4-independent. This deletion was suppressed by processive 5′ end resection and required the 3′>5′ exonuclease activity of the replicative DNA polymerase 8.
- Mre11 nuclease activity suppressed NHEJ and additionally affected the spectrum of events but had no effect on MMEJ.
- Pol ζ and Pol 11 altered NHEJ outcomes in both the presence and absence of Pol4.
- Absence of the Rev3 or Rev7 component of Polζ had different effects on DSB repair. Of particular note, the suppressive effect of Rev7 on MMEJ is consistent with a modulation of resection.
- Long single-strand tails created by resection must be removed to complete MMEJ, but neither Rad1, Mus81 nor Sxl4 was required for this step.
- NHEJ was increased and MMEJ was decreased when ZFN cleavage occurred in non-dividing cells.

## ACKNOWLEDGEMENTS

This work was supported by National Institutes of Health grant R35GM118077 to SJR.

## FIGURE LEGENDS

**Table S1.**
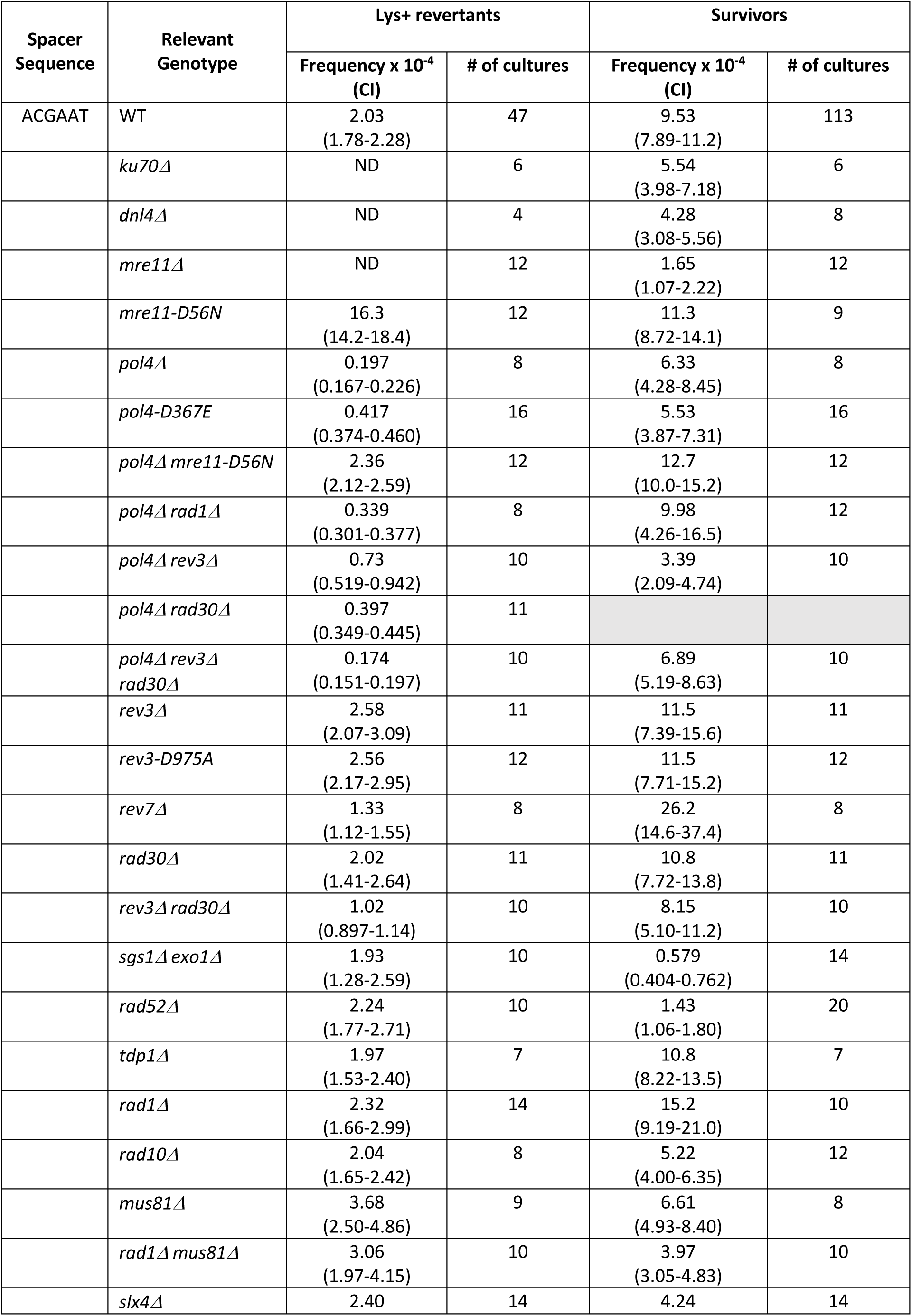
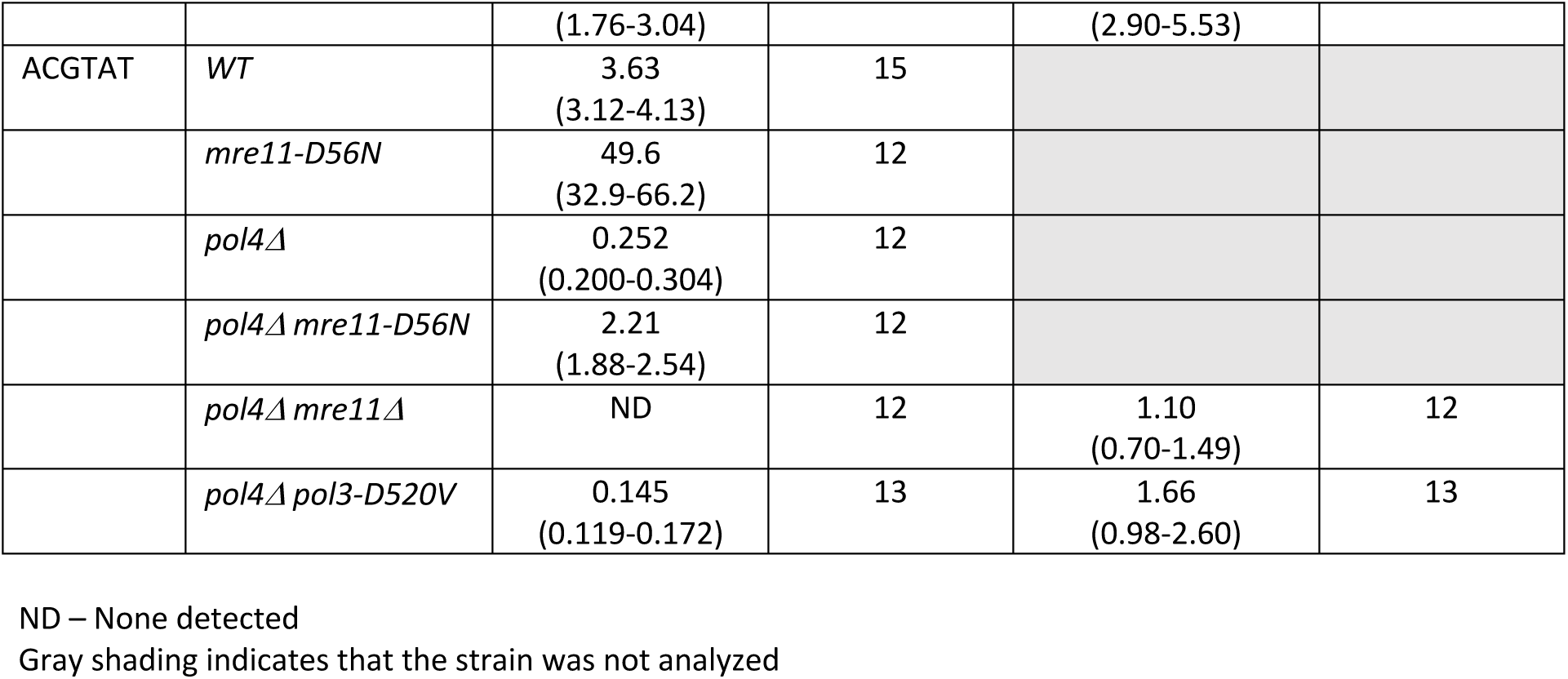
Frequencies of Lys^+^ revertants and of survivors that lost the ZFN cleavage site

**Table S2.**
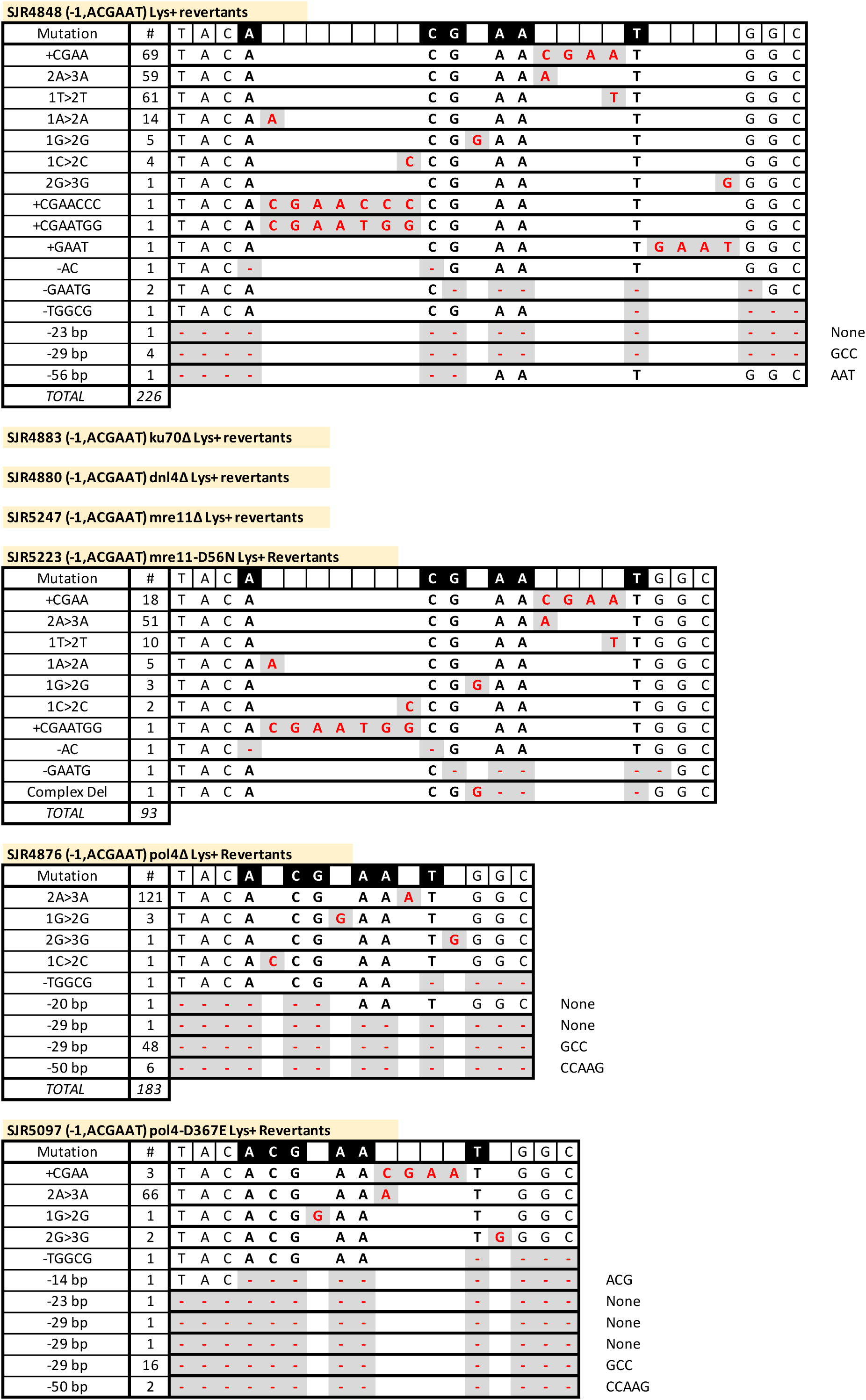
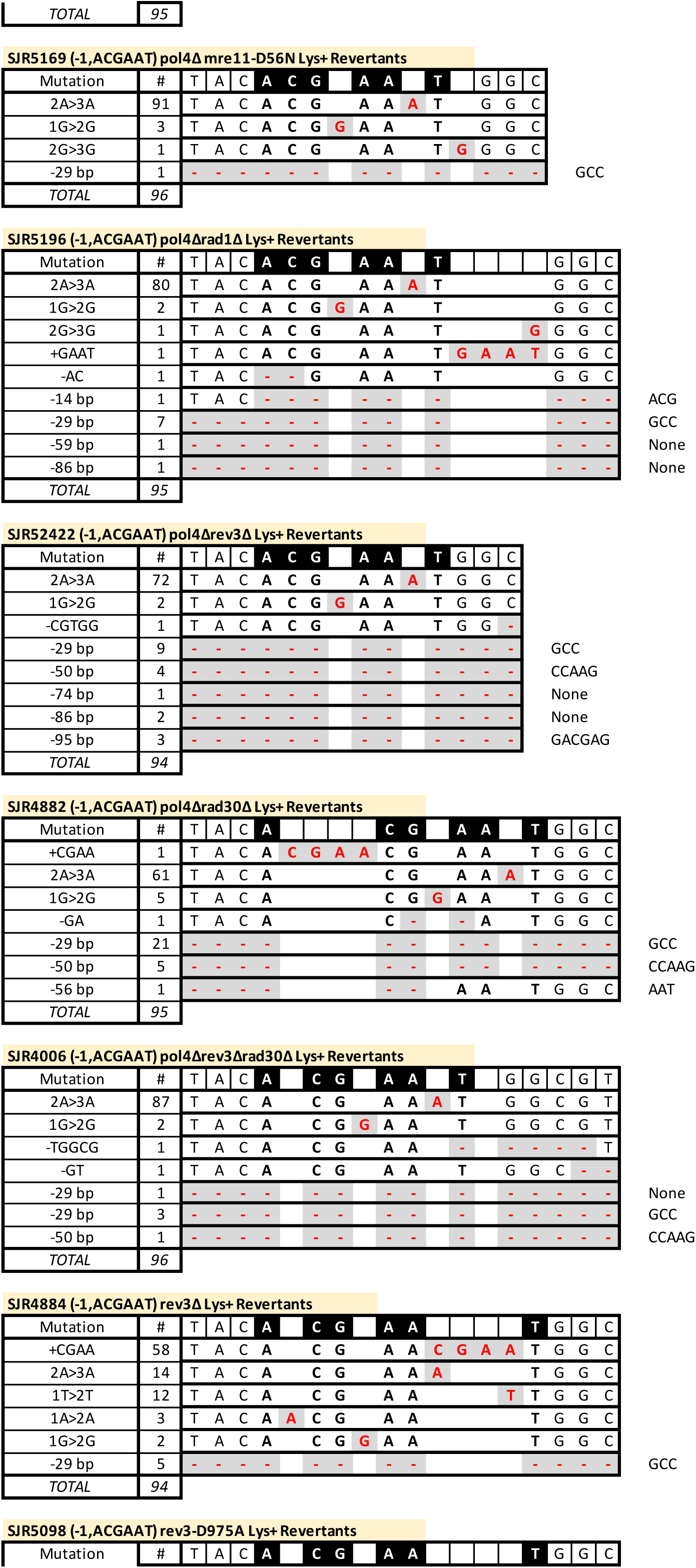
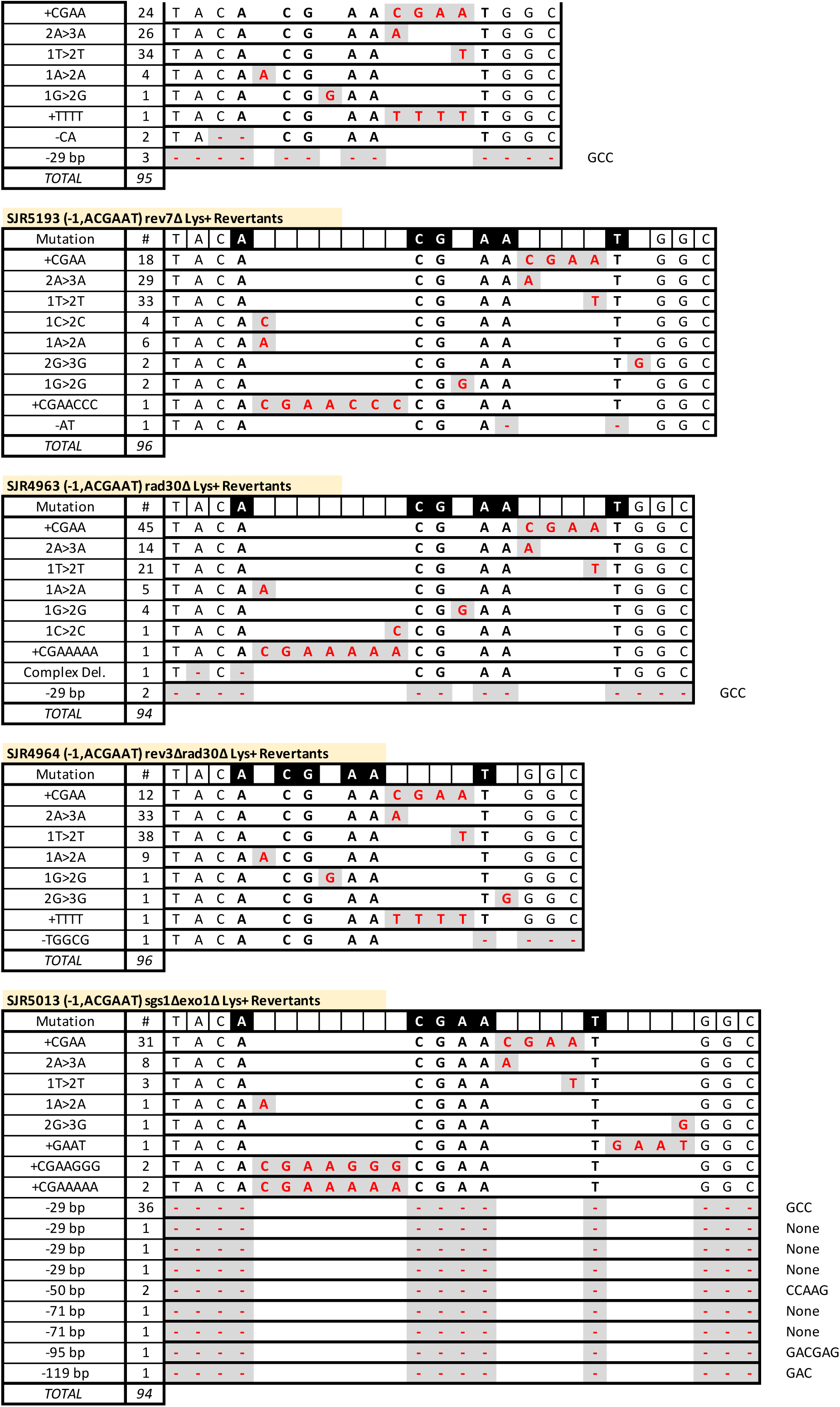
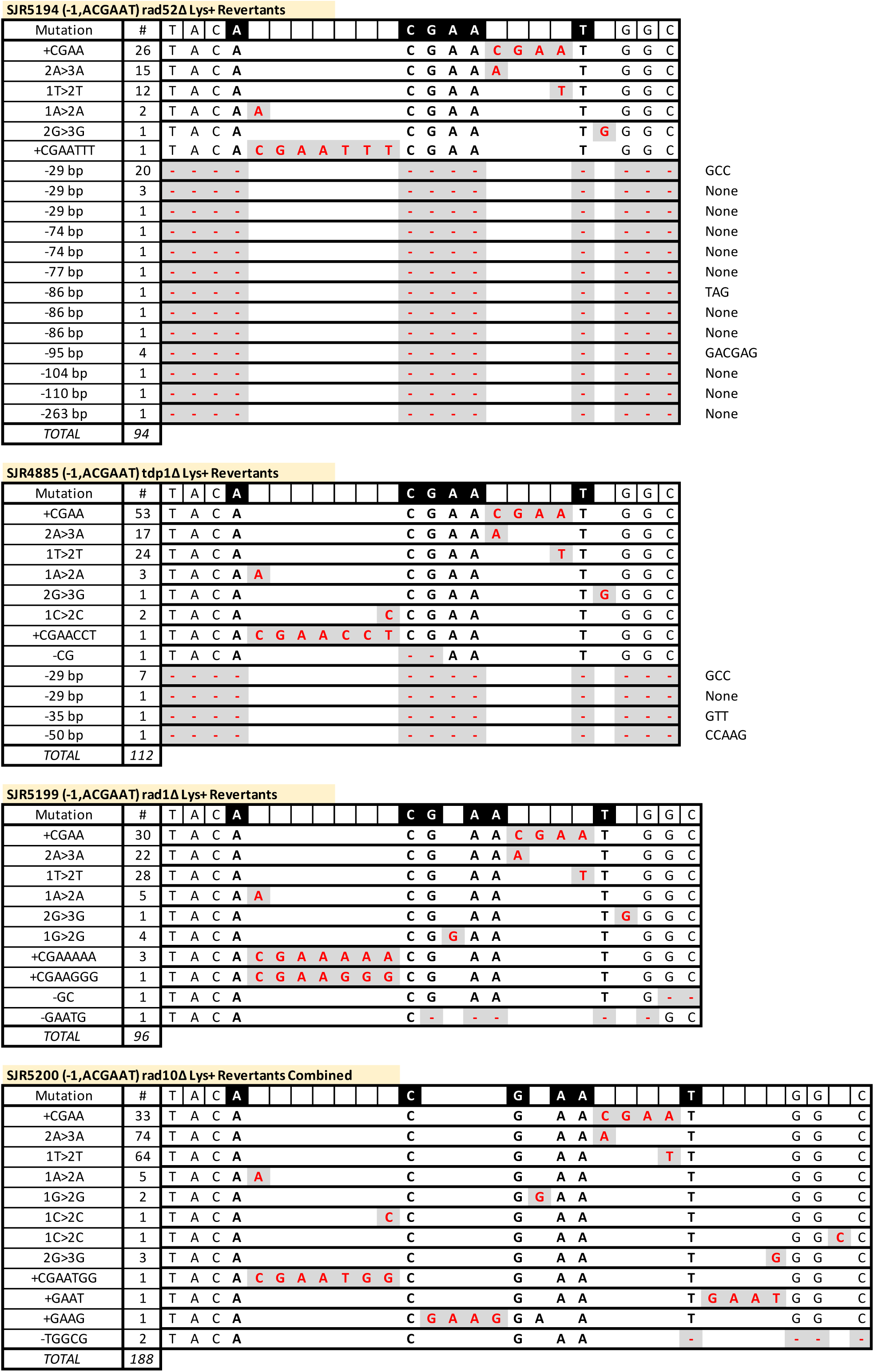
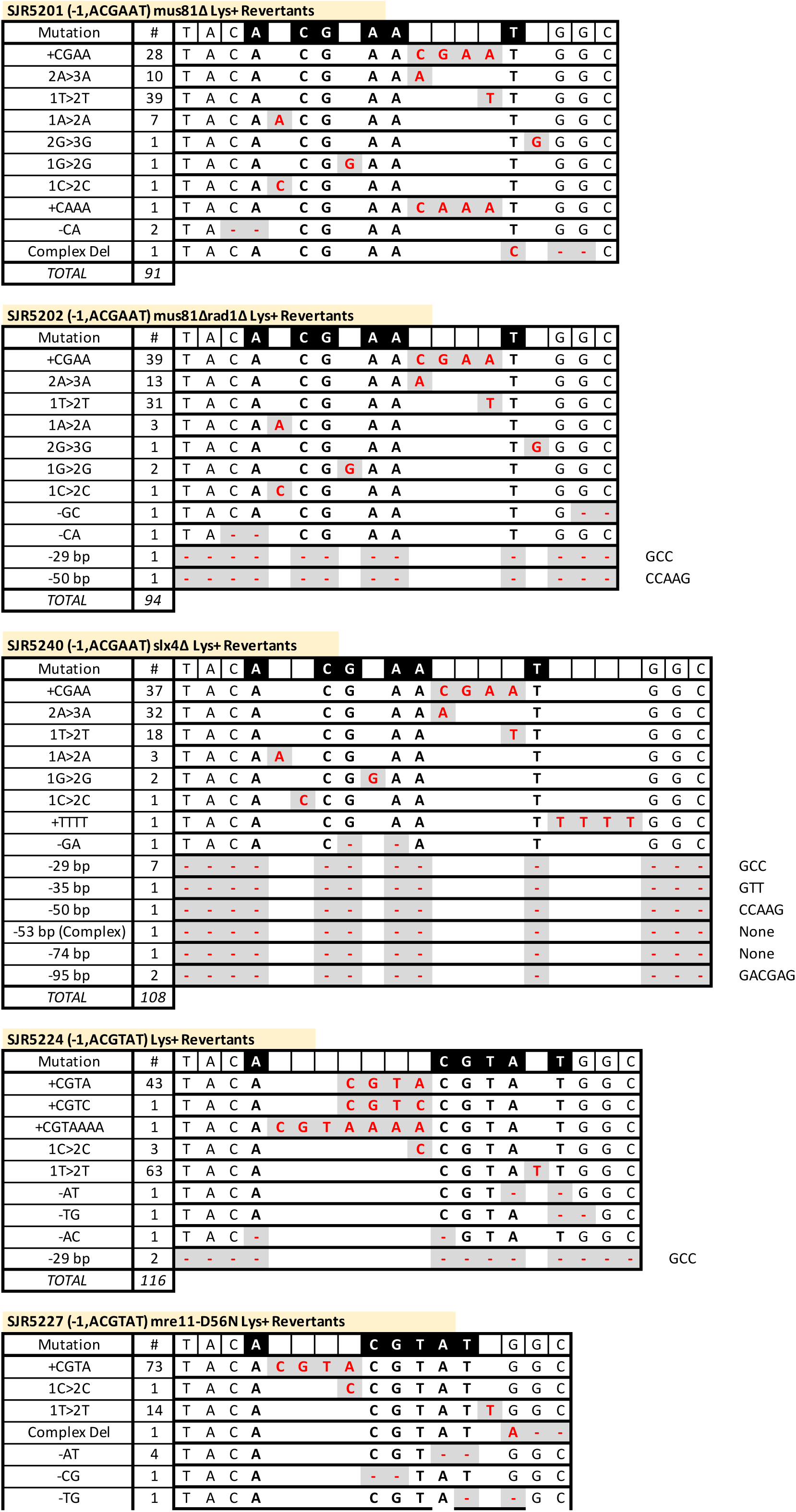
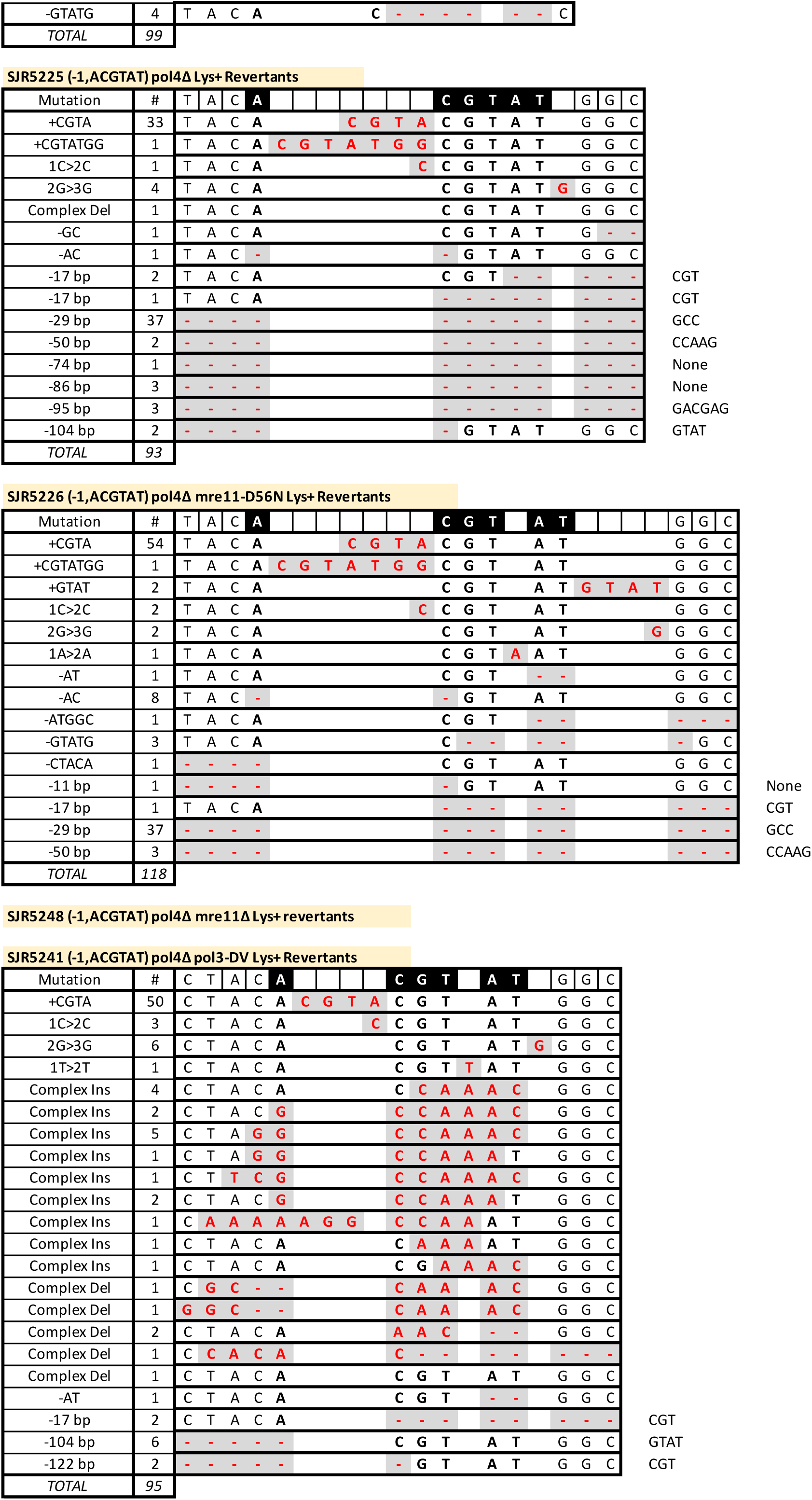
Spectra in Lys+ revertants

**Table S3.**
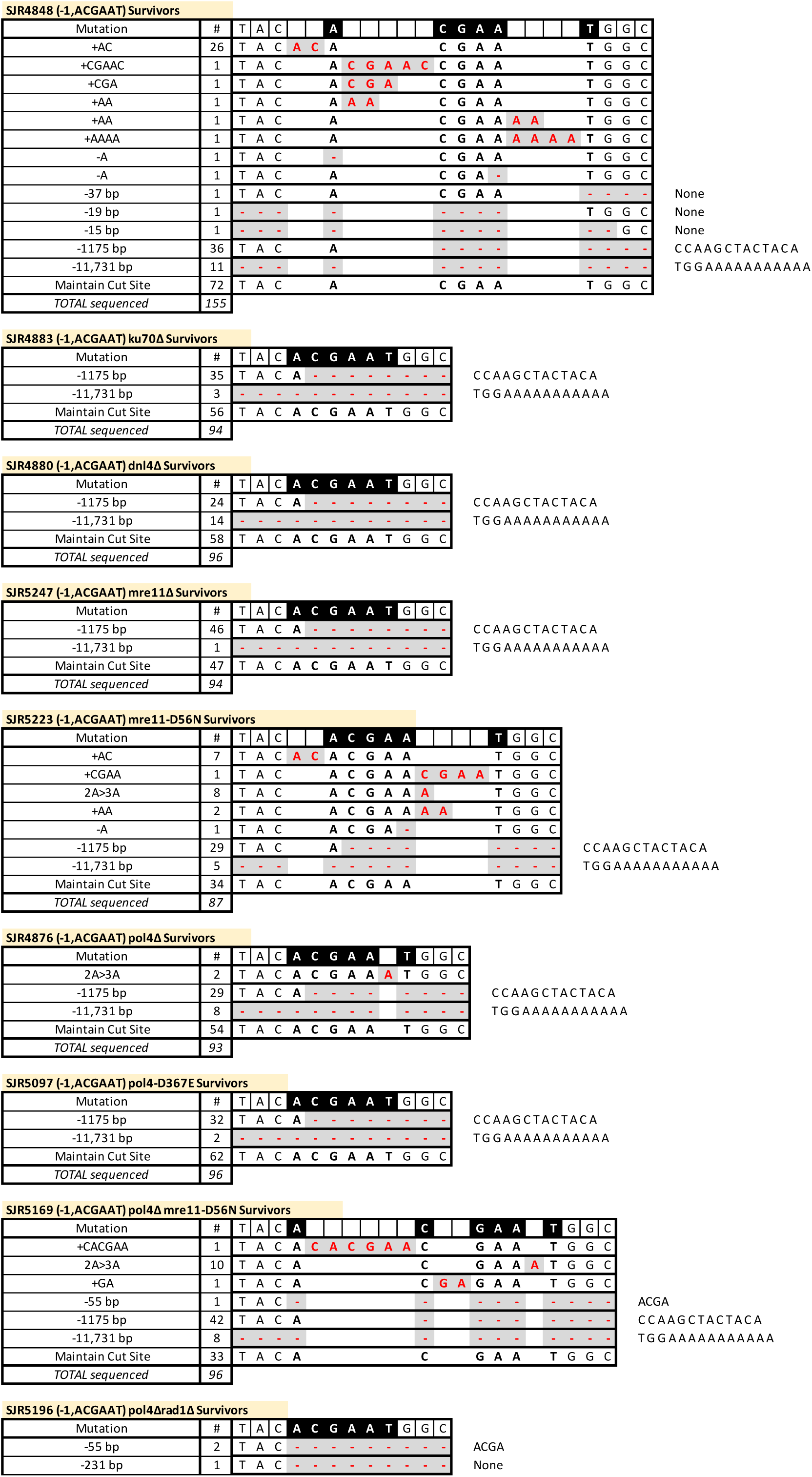
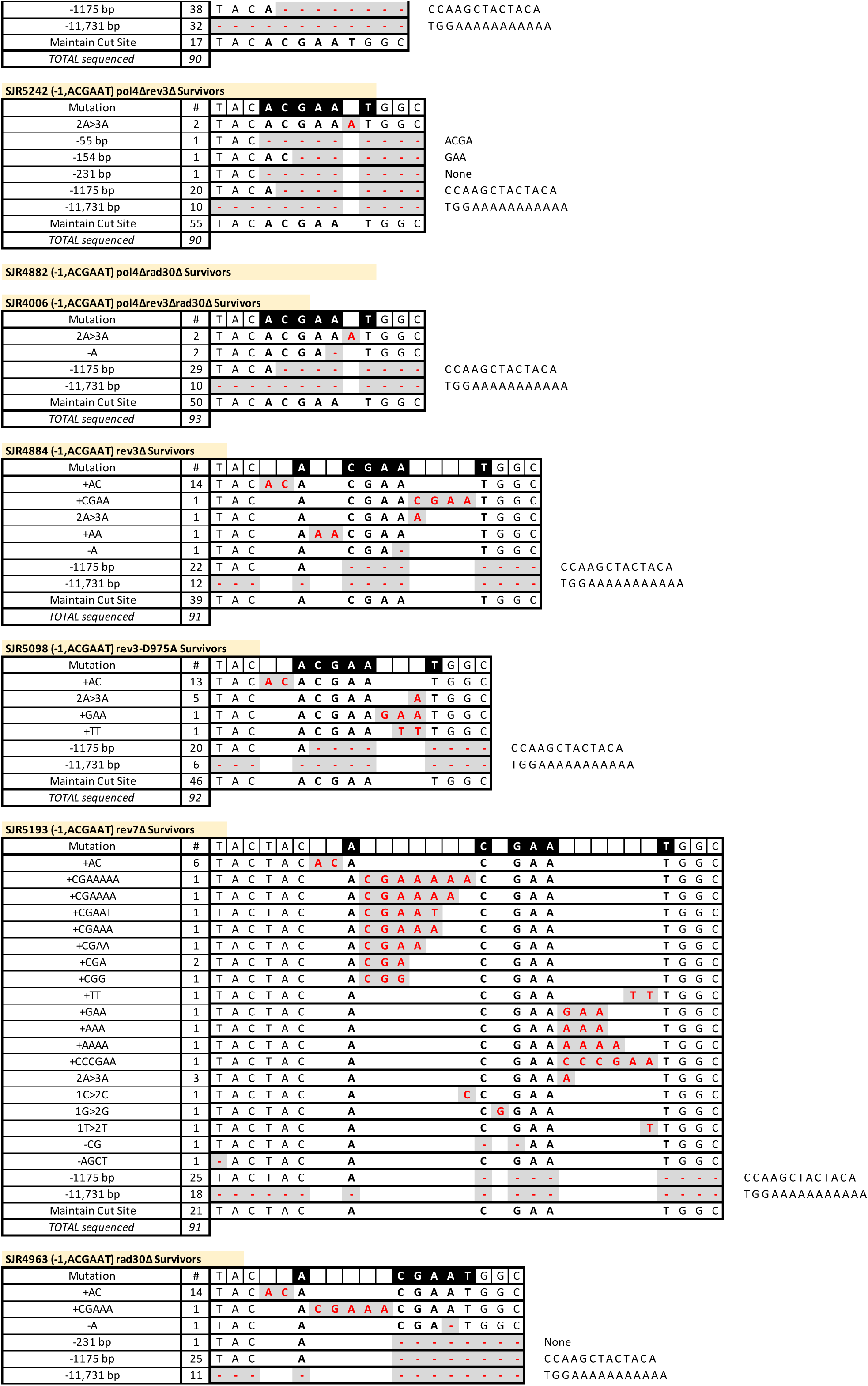
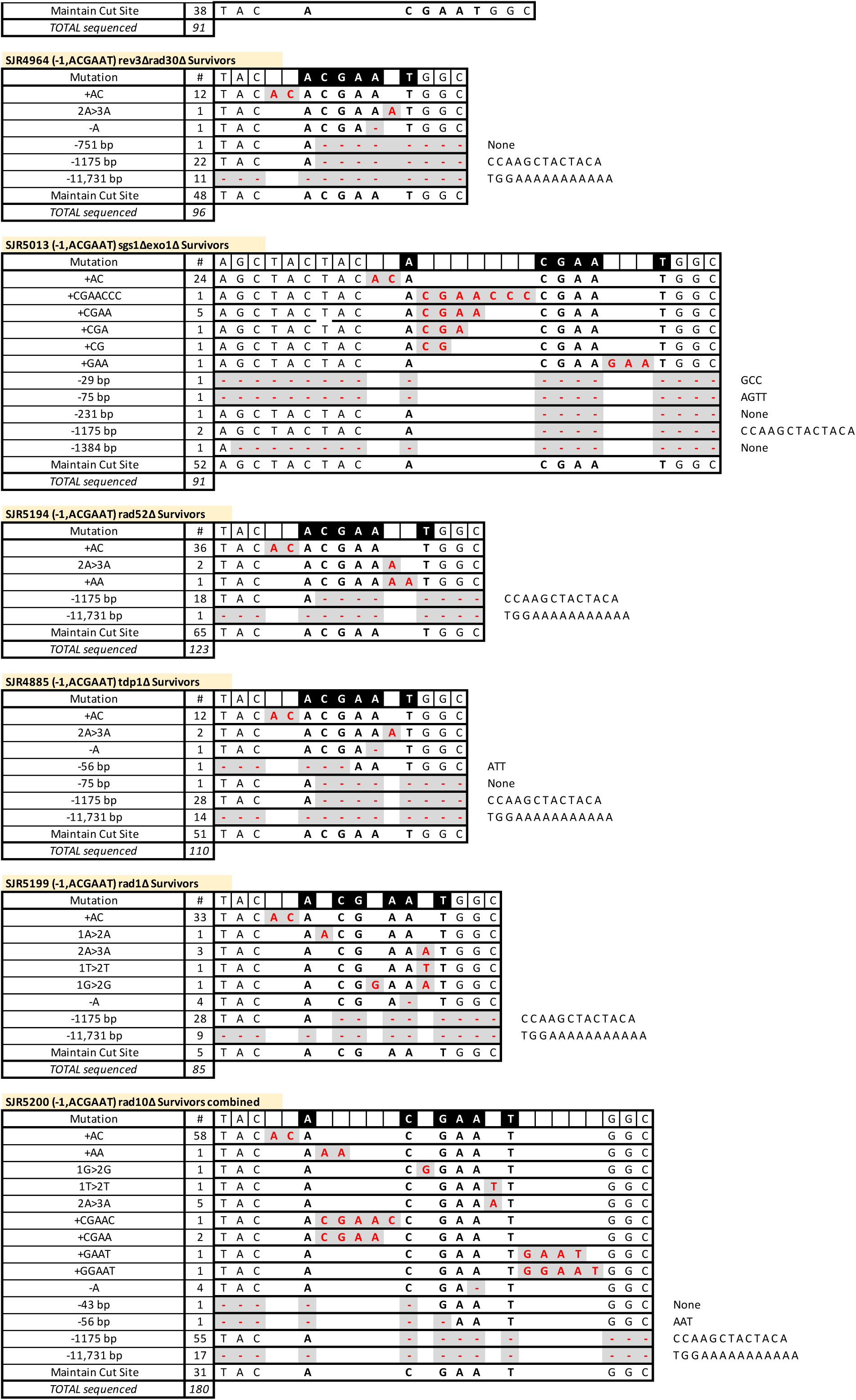
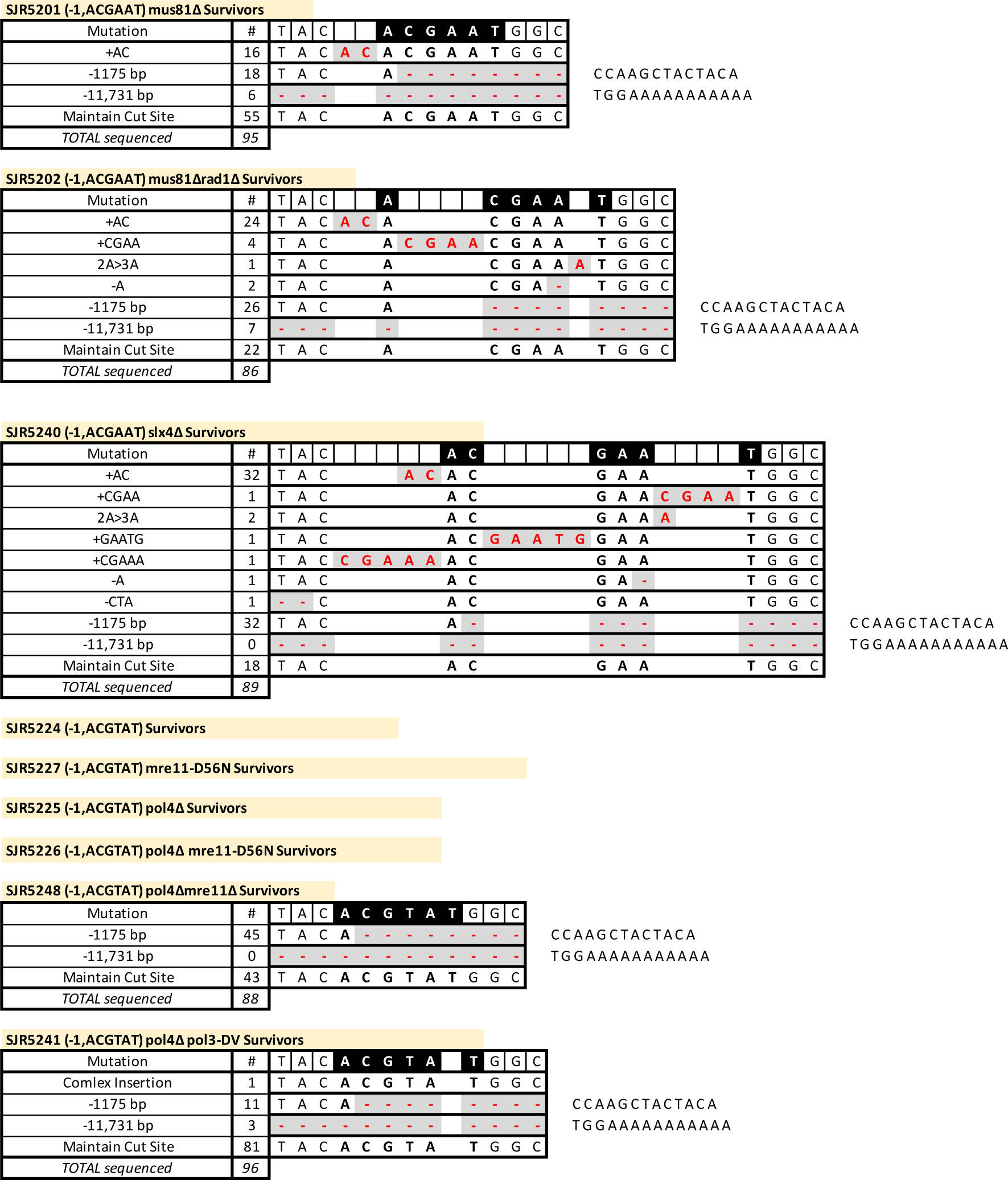
Spectra in survivors

